# Cytoplasmic labile iron accumulates in aging stem cells perturbing a key rheostat for identity control

**DOI:** 10.1101/2021.08.03.454947

**Authors:** Yun-Ruei Kao, Jiahao Chen, Rajni Kumari, Madhuri Tatiparthy, Yuhong Ma, Maria M. Aivalioti, Aliona Zintiridou, Victor Thiruthuvanathan, Julie A. Reisz, Stephanie Stranski, Simone Sidoli, Ulrich Steidl, Angelo D’Alessandro, Britta Will

**Affiliations:** Department of Medicine, Albert Einstein College of Medicine, New York, NY, USA; Department of Cell Biology, Albert Einstein College of Medicine, New York, NY, USA; Department of Biochemistry and Molecular Genetics, University of Colorado Denver – Anschutz Medical Campus, Denver, CO, USA; Department of Biochemistry, Albert Einstein College of Medicine, New York, NY, USA; Ruth L. and David S. Gottesman Institute for Stem Cell Research and Regenerative Medicine, Albert Einstein College of Medicine, New York, NY, USA; Blood Cancer Institute, Albert Einstein Cancer Center, Albert Einstein College of Medicine, New York, NY, USA; Institute for Aging Studies, Albert Einstein College of Medicine, New York, NY, USA

**Keywords:** Adult stem cells, ageing, iron homeostasis, lipid metabolism, histone acetylation, gene regulation.

## Abstract

Bone marrow resident and rarely dividing haematopoietic stem cells (HSC) harbour an extensive self-renewal capacity to sustain life-long blood formation;^1–5^ albeit their function declines during ageing.^6, 7^ Various molecular mechanisms confer stem cell identity, ensure long-term maintenance and are known to be deregulated in aged stem cells.^8, 9^ How these programs are coordinated, particularly during cell division, and what triggers their ageing-associated dysfunction has been unknown. Here, we demonstrate that HSC, containing the lowest amount of cytoplasmic chelatable iron (labile iron pool)^10^ among hematopoietic cells, activate a limited iron response during mitosis. Engagement of this iron homeostasis pathway elicits mobilization and β-oxidation of arachidonic acid and enhances stem cell-defining transcriptional programs governed by histone acetyl transferase Tip60/KAT5. We further find an age-associated expansion of the labile iron pool, along with loss of Tip60/KAT5-dependent gene regulation to contribute to the functional decline of ageing HSC, which can be mitigated by iron chelation. Together, our work reveals cytoplasmic redox active iron as a novel rheostat in adult stem cells; it demonstrates a role for the intracellular labile iron pool in coordinating a cascade of molecular events which reinforces HSC identity during cell division and to drive stem cell ageing when perturbed. As loss of iron homeostasis is commonly observed in the elderly, we anticipate these findings to trigger further studies into understanding and therapeutic mitigation of labile iron pool-dependent stem cell dysfunction in a wide range of degenerative and malignant pathologies.

---

HSC, as various other somatic stem cells, reside mostly in a quiescent state^3^; infrequently dividing only when cued by distinct intrinsic^11^ and extrinsic signals^12^ to enter the active cell cycle and undergo cell division to contribute to mature blood cell production or replenish the stem cell pool^13, 14^. Accumulation of genetic and epigenetic gene-regulatory alterations occur in HSC over time and multiple replications^15, 16^. They, along with various extracellular changes^17–19^, contribute to the ageing-related decline of functional stem cells^20^ which are the cellular source of several degenerative^21^ and malignant pathologies^22^.

Past studies have provided crucial insights into the molecular and cellular mechanisms governing long-term maintenance of adult stem cells; these include, ability to undergo and modulate cell division modes (asymmetric vs. symmetric)^23–25^, reliance on glycolytic pathways^26^ and autophagy^9, 27^ for curtailing reactive oxygen species (ROS) generation while producing sufficient energy and cellular building blocks^28, 29^, and low protein translation rates^30–33^. Each cell division demands wide-ranging metabolic and gene-regulatory adaptations to meet the changing energetic and structural demands of mitosis^34–38^. As these adaptations are largely incompatible with their sustained long-term function, stem cells must counteract mitosis-related molecular perturbations - the molecular processes at play have remained elusive thus far.

While ensuring a sufficient amount of redox active, readily available iron which is required in numerous electron transfer reactions governing fundamental cellular processes^39^, cells tightly regulate the size of the intracellular labile iron pool (LIP)^10^ to limit adverse ROS generation^40, 41^. Perturbations in the ability to limit intracellular iron is detrimental for cells^42^ and known to compromise HSC maintenance and function via altered redox signalling and increased macromolecule oxidation and damage^43^. We have previously uncovered that iron chelator exposure increases the number of functional HSC *ex vivo* and *in vivo*^44^. The HSC stimulatory effects of iron chelator (IC) treatment and the well characterized central roles of redox active intracellular iron in sustaining basic cell function prompted us to examine a potential regulatory role of the LIP in controlling somatic stem cell function.

## HSC contain the most limited LIP among hematopoietic cells

The size of readily accessible functional intracellular iron pools is determined by the cellular state and function of cells^45, 46^. We first quantified the amount of intracellular chelatable, redox active ferrous iron (Fe^2+^) species across various haematopoietic cell types which revealed differences in their LIP. Compared to highly proliferative Lineage^-^ (Lin^-^) and myeloid progenitor cells (Lin^-^cKit^+^Sca-1^-^, LK), most immature hematopoietic stem cells (Lineage^-^ckit^+^Sca-1^+^CD48^-^CD150^+^^47^ or CD34^-^HSC^48^, HSC) and multipotent progenitor cells (Lineage^-^ckit^+^Sca-1^+^CD48^-^CD150^-^^47^, MPP) harbour well detectable, but significantly lower amounts of free Fe^2+^ (**Extended Data Fig. 1a**), consistent with their mostly quiescent state and lower energetic requirements^26, 49–52^.

**Figure 1:**
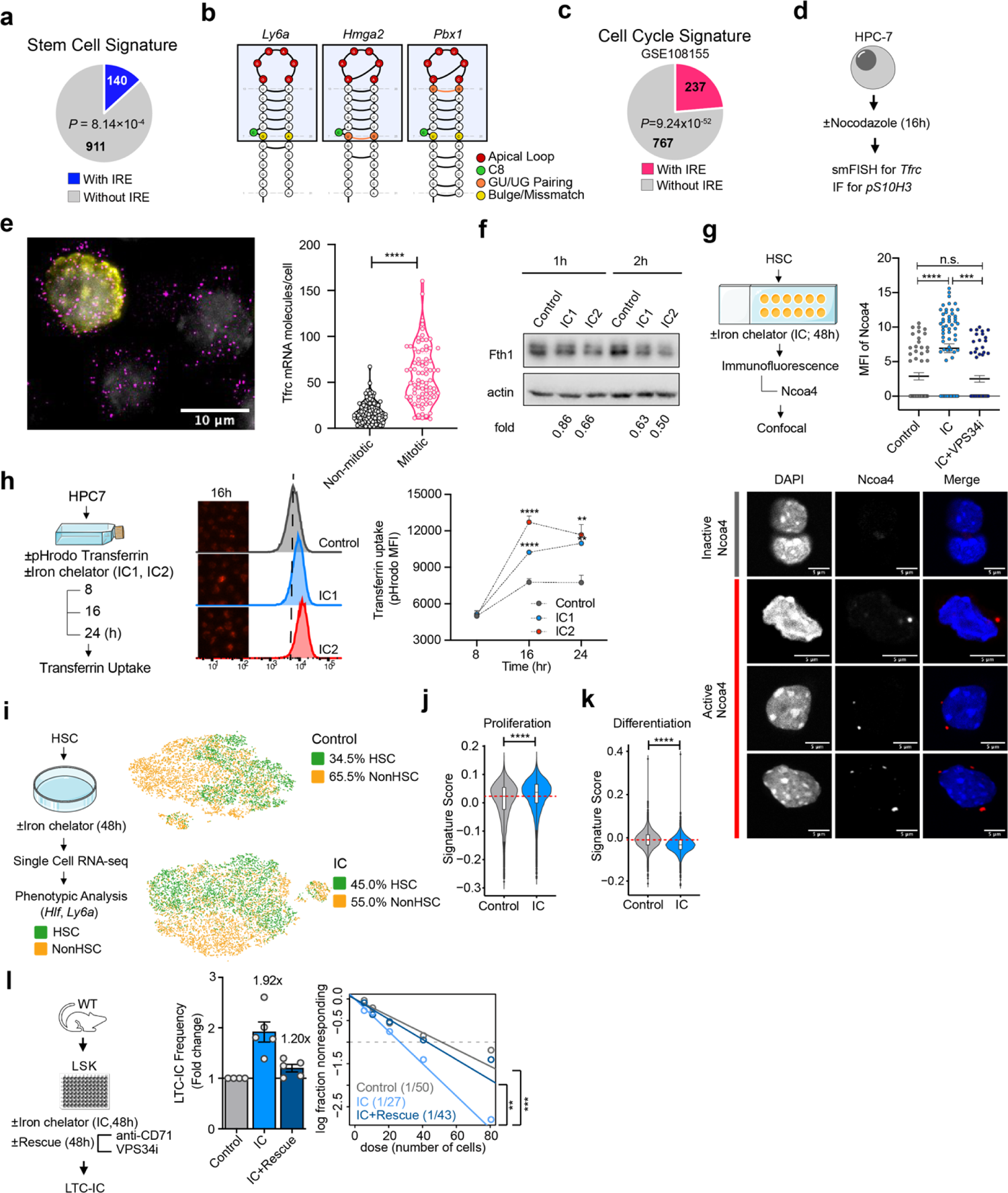
Activation of the limited iron response increases self-renewal of multipotent HSPC. **a,b,** Quantification of iron responsive elements (IRE) in mRNA transcripts of HSC gene signatures; **a,** IRE enrichment analysis of HSC gene signatures; **b,** IRE-motif in *Ly6a*, *Hmga2*, and *Pbx1*. **c**, IRE enrichment analysis of genes associated with HSC cycling. Cell cycle gene signatures were defined with DEG in Retinoic Acid reporter positive (highly proliferative) compared to negative (less proliferative) mouse HSC (GSE108155). **d-e,** Scheme showing experimental design: HPC-7 cells were cultured in the presence of vehicle control (0.01%DMSO) or Nocodazole for 16hrs and subjected to sequential *Tfrc* smRNA FISH followed by phosphoSer10 Histone H3 (pS10H3) immunofluorescence to identify mitotic cells (**d**). **e**, Left: Representative filtered and overlaid images of treated HPC-7 cells stained by *Tfrc* smRNA FISH (Cy5, pseudocolor magenta), phosphoSer10 Histone H3 (AlexaFluor 488, pseudocolor yellow) and nuclei (DAPI, pseudocolor gray). Scale bars, 10 μm; Right: Cells positive for pS10H3 staining are marked as mitotic (n=80) and cells negative for pS10H3 staining are marked as non-mitotic (n=104). Violin plots show absolute number of cytoplasmic *Tfrc* mRNA molecules per cell. **f-h**, Activation of iron homeostasis pathways upon iron chelator (IC1: 5µM DFO; IC2: 3µg/ml EP) exposure as indicated by rapid degradation of ferritin heavy chain (Fth1) protein in HPC-7 cells (**f**); Ferritinophagy in HSC mediated by Naco4 activation detected by immunohistochemistry staining (**g**), n = 55 (Control), 66 (IC), 62 (IC+VPS34i) cells; and transferrin uptake in HPC-7 cells (**h**), n = 9; 2 independent experiments. **i,** Quantification of phenotypical HSC by scRNA sequencing after 48hrs IC (DFO) exposure (mock-treated HSC as controls). Cells with *Hlf* and *Ly6a* expression higher than the 25% percentile of their average expression in all cells in both IC and control groups were defined as HSC (green), otherwise non-HSC (orange). **j-k,** Comparison of the expression of IC-treated HSC versus control HSC in scRNA RNA sequencing, with gene signatures associated with HSC proliferation (**j,** GSE108155) and differentiation (**k**). Lists of the gene signatures are available in **Extended Data Table 1**. Signature scores were calculated with *Seurat* package. **l,** Long-term culture-initiating cell (LTC-IC) assay to quantify HSC upon IC exposure alone or along with inhibition of iron homeostasis pathways. Left: Experimental strategy for pharmacological inhibition of iron import and intracellular mobilization. Right: Fold changes of long-term culture-initiating cell (LTC-IC) frequencies compared to controls across individual mice (bar graph). n = 5. LTC-IC frequencies was estimated by ELDA and shown in parentheses in ELDA plot (right). If not specified otherwise, data are mean ± SEM (**e**, **g, h, l** (bar graph)). Significance *p*-values, indicated as *p < 0.05, **p < 0.01, ***p < 0.001, ****p < 0.0001, n.s. (not significant), were calculated using Student’s *t*-test (unpaired: **e**, **g, j, k** (bar graph); paired: **h**), Poisson statistics (**l** (ELDA plots)). Significance of enrichment was calculated by hypergeometric test (**a, c**).

Several key regulators for iron sensing, transport and storage are well detectable and at comparable levels in mouse HSC and maturing progenitor cell populations (**Extended Data Fig. 1b-d**); notable exceptions are iron storage genes *Ferritin Heavy Chain 1* (*Fth1*) and *Ferritin Light Chain* (*Ftl1*) which are found expressed at higher levels in HSC than in more mature progenitor cells, while iron sensors *Aconitase 1 / Iron responsive protein 1* (*Aco1/Irp1*) and *Iron Responsive Element Binding Protein 2* (*Ireb2*) and the *Transferrin Receptor* (*Tfrc*), relevant for cellular iron uptake, showed slightly lower abundance in HSC compared to their mature progeny (**Extended Data Fig. 1b-d**). Quantification of Transferrin Receptor protein (CD71) expression and cell surface presentation further showed that HSC harboured an approximately 70% lower total protein content than erythro-myeloid lineage-committed progenitor cells, in line with previous observations^25, 53^; notably, our analyses also revealed that only a minor fraction of the receptor is presented on the surface of all cell types tested (**Extended Data Fig. 1e,f**). These data support a lower labile iron requirement and a bias towards iron storage in HSC under steady state conditions.

## Limited iron response activation in mitotic stem cells

A decrease in intracellular labile iron content is sensed and corrected by the iron responsive protein (IRP) / iron responsive element (IRE) system which post-transcriptionally controls the expression and translation of several mRNA transcripts encoding proteins of iron-, oxygen- and energy metabolism, including the stabilization of *Tfrc* transcripts (limited iron response)^54^. Motif prediction in mouse and human HSC gene expression signatures showed a significant enrichment of IRE bearing transcripts to a similar extent as found when using gene expression signatures of erythroid cells whose function critically relies on iron^55–57^ (**Fig. 1a,b, Extended Data Fig. 1g-j**). We noted that almost a quarter of differentially expressed transcripts in proliferating HSC (vs. quiescent stem cells) contained IRE motifs (**Fig. 1c, Extended Data Fig. 1k**), suggesting an activation of the IRP/IRE-mediated limited iron response in proliferating HSC. To test this prediction, we quantified cytoplasmic and nuclear *Tfrc* transcript levels, as well as transcription dynamics at single cell and single-molecule resolution. We utilized single-molecule RNA fluorescence in situ hybridization (smRNA FISH) technology, which we have reported and validated in the hematopoietic system previously^58^, in a mouse HSC cell line, HPC-7 (**Fig. 1d, Extended Data Fig. 2a-g**). Compared to non-mitotic cells, mitotic HSC showed a significant increase in cytoplasmic mRNA levels (**Fig. 1e, Extended Data Fig. 2h**) while displaying no signs of increased transcriptional activity at the *Tfrc* locus as evidenced by unchanged transcriptional burst size and frequency (**Extended Data Fig. 2i,j**). Experimental decrease of intracellular iron through exposure to an IC (deferoxamine (DFO)) showed similar results (**Extended Data Fig. 2k-n**). These observations revealed that the increased abundance of *Tfrc* is mediated through enhanced mRNA stabilization^54^ and strongly suggested that iron limitation triggers a compensatory iron homeostasis reaction in dividing HSC.

**Figure 2:**
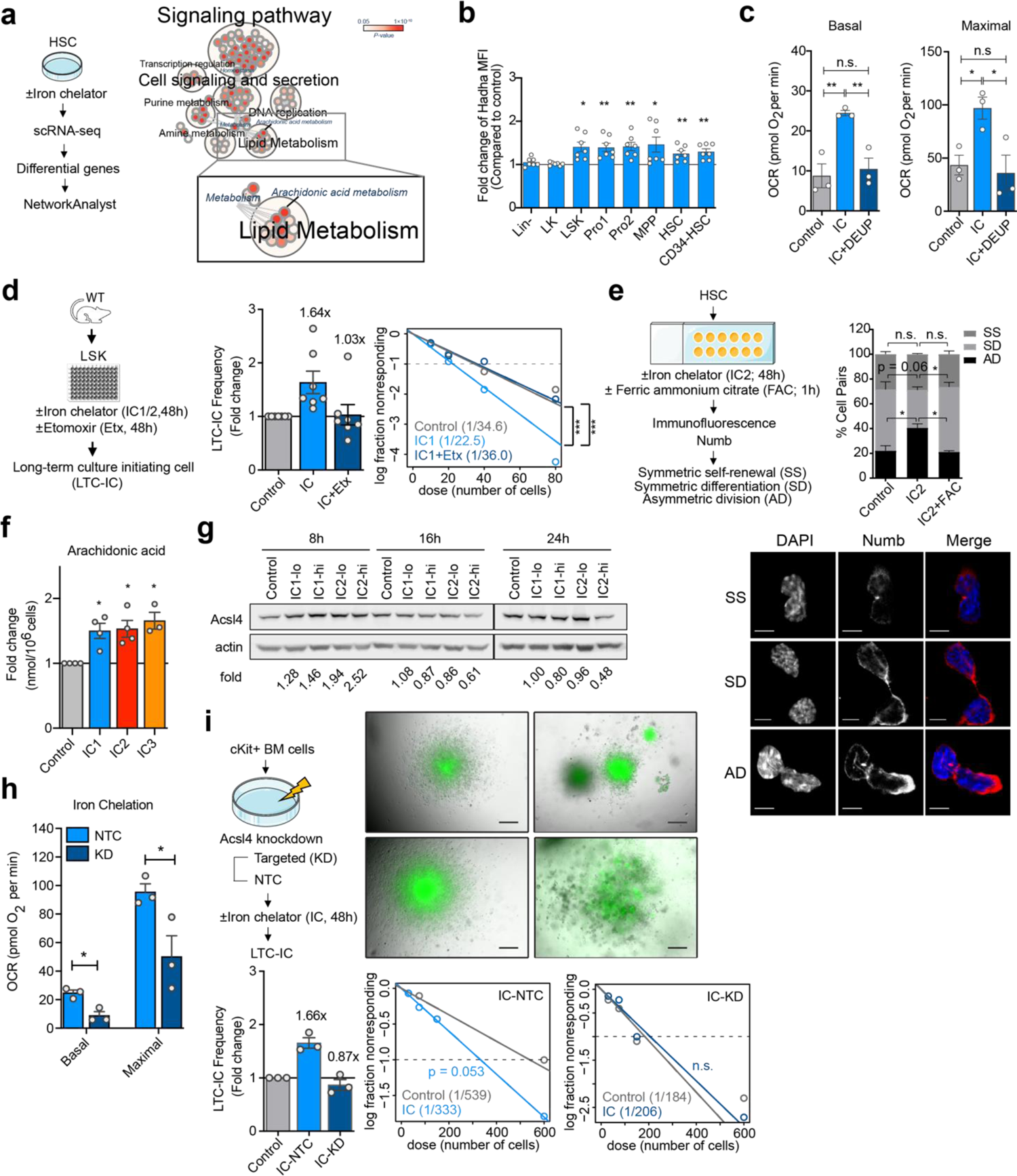
Limited iron response-triggered β-oxidation of arachidonic acid increases HSPC function. **a,** Pathway network analysis of molecular alterations in HSC upon IC exposure. Left: Following IC (DFO) treatment, HSC were subjected to scRNA-seq and DEG were subjected to pathway analysis by *NetworkAnalyst*; Right: Significantly enriched pathways (*p* < 0.05) were subjected to *EnrichmentMap* for network visualization, and clustered with *AutoAnnotate* based on the coefficient calculated by the overlapping of implicating genes. Each circle represents a pathway, the color intensity is a mapping of enrichment significance *p*-value. Thickness of connecting lines represents the similarity coefficient calculated by *EnrichmentMap*. **b,** Quantification of Hadha levels in different hematopoietic stem and progenitor cell populations 24hrs after IC (DFO) exposure compared to vehicle controls. n = 7. **c,** Assessment of the effect of iron chelation on fatty acid metabolism in Ba/F3 cells, by Seahorse analysis detecting β-oxidation (after CPT-1 inhibition with etomoxir) specific oxygen consumption rates (OCR). Endogenous basal (left) and maximal (right) fatty acid oxidation rates are shown. See methods for details in the calculation of β-oxidation specific OCR. **d,** LTC-IC assay to quantify functional HSC upon IC1 (DFO) treatment alone or in combination with pharmacological inhibition of fatty acid import to mitochondria using etomoxir (Etx). Bar graphs represent foldchanges of LTC-IC frequency of chelator treatment compared to control for individual mice. In the ELDA plot, average stem cell frequency is shown for each treatment group. n = 7. **e,** Quantification of symmetric self-renewal (SS), symmetric differentiation (SD), and asymmetric cell divisions (AD) based on Numb protein (red) abundance. Nucleus was counterstained with DAPI (blue), scale bar indicates 5 µm. Bar plot shows the quantification of division modes of HSC treated with IC2 (EP) alone or in combination with ferric ammonium citrate (FAC). Shown is the percentage of HSC daughter cell pairs (20-30 cell pairs scored per condition and experiment) for each division mode. n = 2 independent experiments. **f,** Quantification of AA levels by mass spectrometry following IC treatment (IC1: DFO, IC2: EP, IC3: DFX) of HPC-7 cells. n = 3-4. **g,** Quantification of Acsl4 protein levels by Western blot in HPC-7 cells after IC exposure (IC1: DFO, IC2: EP) at the time point of 8, 16 and 24hrs. **h,** Assessment of fatty acid oxidation rates in BA/F3 cells transduced with non-targeting control (NTC) or shRNA targeting *Acsl4* (KD) after IC (DFO) treatment. Basal and maximal fatty acid-specific OCR are shown. n = 3. **i,** HSC enumeration by LTC-IC following RNAi-mediated ablation of *Acsl4* (KD) and IC (DFO) treatment; assessment of shRNA construct vs. a non-targeting control (NTC) shRNA vector. Representative of LTC-IC GFP+ colonies are shown. Bar plot shows the quantification of LTC-IC frequency estimated in ELDA plots, n = 3. If not specified otherwise, data are mean ± SEM (**b, c, e, f, h; d and i** (bar graph)). Significance *p*-values, indicated as *p < 0.05, **p < 0.01, n.s. (not significant), were calculated using paired Student’s *t*-test (**b, c, e, f**, and **h**), or Poisson statistics (**d, i** (ELDA plots)).

We next delineated the molecular response of HSC to acute iron limitation in detail for which we employed short term *ex vivo* IC treatment of primary HSC and HPC-7 cells. Compared to vehicle treated controls, IC exposure to a rapid decrease in Fth1 and Ftl protein abundance (**Fig. 1f, Extended Data Fig. 3a**) within one hour, activation of intracellular iron storage mobilization by Ncoa4-dependent ferritinophagy^59^ (**Fig. 1g, Extended Data Fig. 3b**) and was associated with elevated and readily restored intracellular Fe^2+^ levels within only four hours of IC exposure (**Extended Data Fig. 3c**). We also found alterations in mRNA expression consistent with reduced intracellular iron content after IC exposure of primary HSC for 16 hrs (**Extended Data Fig. 3d-f, Extended Data Table 2**). Notably, stem cells did not show a detectable increase in the uptake of extracellular iron before 8 hrs (**Fig. 1h**) or elevated CD71 cell surface presentation before 6 hrs (**Extended Data Fig. 3g**) after IC treatment. Yet, we also noted that intracellular iron stores seem smaller in hematopoietic stem and multipotent stem cells as compared to their more mature progeny (**Extended Data Fig. 3h**). This set of data strongly suggests that upon activation of the limited iron response HSC, first increase intracellular iron mobilization before upregulating CD71 which has previously been associated with enhanced differentiation commitment^25, 53^.

**Figure 3:**
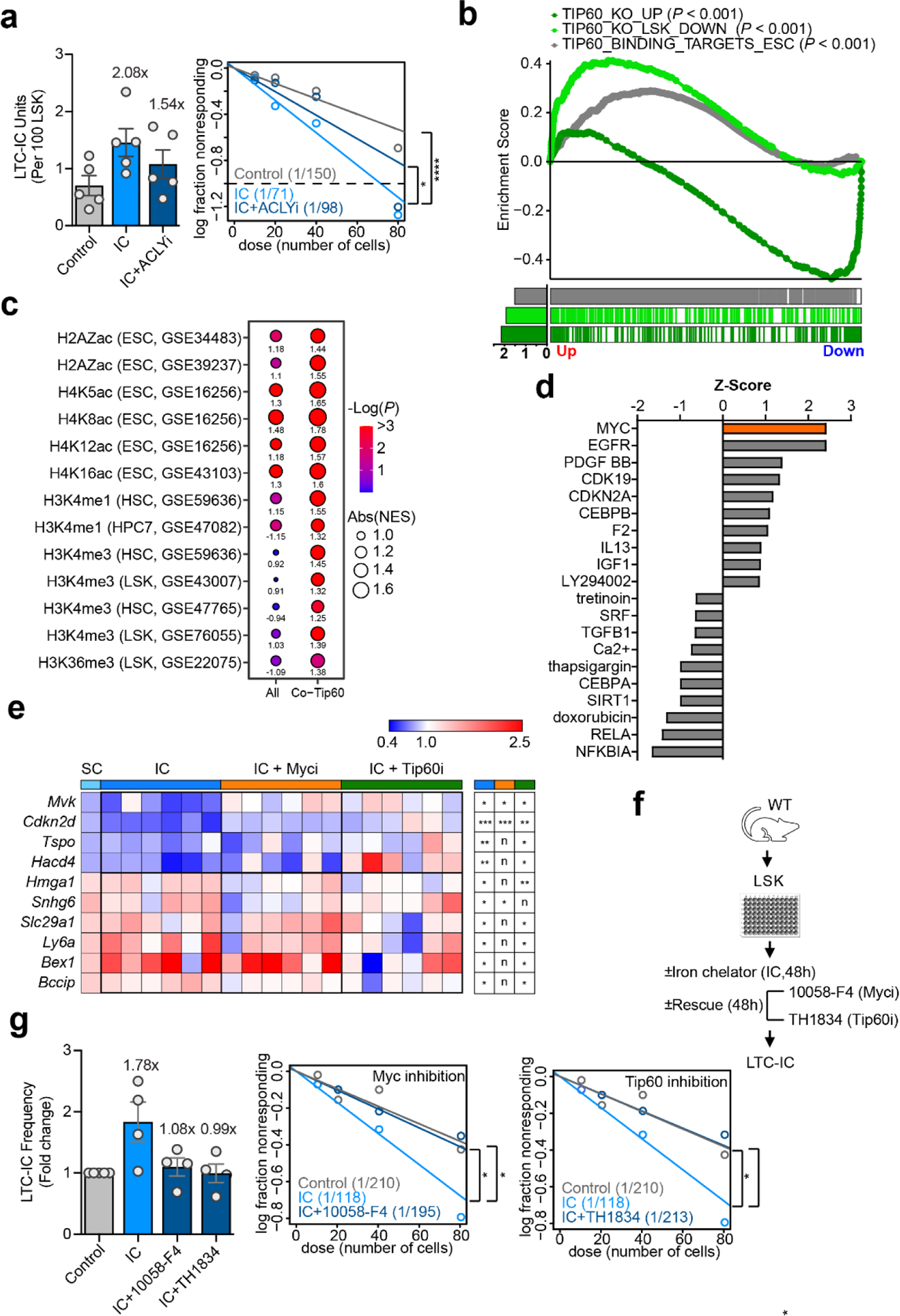
Tip60 controls HSC-typic gene expression upon activation of the limited iron response. **a,** Quantification of HSC frequency by LTC-IC assay after IC treatment alone or in combination with ATP citrate lyase inhibition (ACLYi). n = 4. Bar graphs represent foldchanges of LTC-IC frequency of chelator treatment compared to control for each individual mouse. Average LTC-IC frequencies are shown in parentheses. Significance tested using ELDA, \**p* < 0.05. **b,** *GSEAPreranked* enrichment analysis of scRNA-seq data using gene set of Tip60 binding targets in ESC, or DEG in Tip60-KO LSK. Bar plot shows the normalized enrichment score (NES, absolute value) in the respective gene set. **c,** Balloon plot showing *GSEAPreranked* enrichment analysis performed with gene sets associated with histone modifications and Tip60 binding. NES and significance *p*-values of gene sets with all the genes associated with each of the histone modifications and those common with Tip60 binding targets are shown. **d,** Upstream regulator analysis of differentially expressed genes upon IC exposure of HSC that are also direct targets of Tip60. **e,** Expression changes of Tip60/Myc target genes in HSC after IC treatment alone or in combination with inhibition of c-Myc (Myci) or Tip60 (Tip60i) by Fluidigm analysis. Fold changes of genes across treatment groups from Fluidigm and scRNA-seq (SC) are shown. Significance of differential expression shown (right panel) were estimated by paired Student’s *t*-test comparing the Δ^Ct^ in IC versus control, or co-treatment versus IC. *p < 0.05, **p < 0.01, ***p < 0.001, *n*, not significant. **f-g,** Quantification of HSC by LTC-IC assay after IC treatment alone or in combination with Tip60, or Myc inhibition. n = 4. Average LTC-IC frequencies are shown in parentheses. Significance tested using ELDA, \**p* < 0.05.

## Activation of the limited iron response increases HSC function

To test whether alterations in LIP size and activation of the limited iron response has functional consequences in HSC, we exposed highly purified stem cells to IC *ex vivo* for 48 hrs and subsequently preformed single cell transcriptomic analysis. Compared to vehicle treated control HSC cultures, IC treated cells showed a larger proportion of cells retaining HSC-typic gene expression signatures (**Fig. 1i, Extended Data Fig. 4a**), with increased scores of gene signatures for *HSC proliferation*, and *Stem Cell identity*, as well as concomitantly decreased scores for *HSC differentiation* and *quiescence* (**Fig. 1j,k, Extended Data Fig. 4b-d, Extended Data Table 1**). We next subjected mice to a 14-day regimen of daily treatment with iron chelator DFO^60, 61^. Compared to mock-treated controls, this treatment regimen led to reduced body iron without triggering overt iron deficiency pathology (**Extended Data Fig. 4e-g**). Hematopoietic stem and multipotent progenitor cell (HSPC) activity, assessed by long-term culture-initiating cell (LTC-IC) quantification, was found increased in IC-treated mice compared to mock control animals (**Extended Data Fig. 4h**), confirming our previous observations with human HSPC and additional iron chleators^44^. These findings indicated that IC exposed cells retained more effectively their stem cell-typic gene expression programs and function compared to vehicle controls.

**Figure 4:**
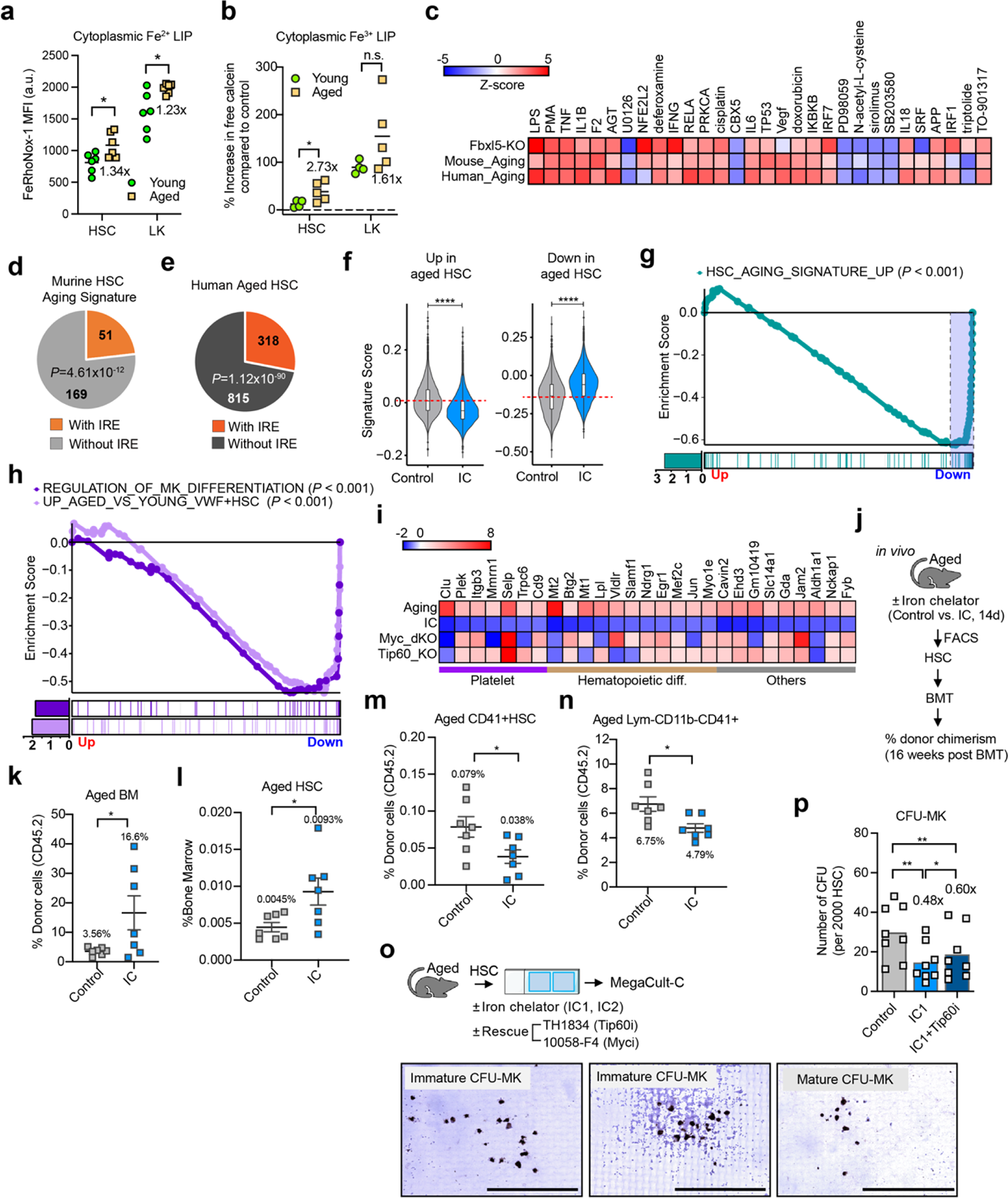
Ageing associated cytoplasmic iron loading impairs HSPC function. **a,b,** Increased labile iron pool (LIP) in aged HSC compared to young HSC. Cytoplasmic Fe^2+^ (**a**) or Fe^3+^ (**b**) LIP were measured using FeRhoNox-1 or Calcein-AM staining, respectively. For Calcein-AM assay, EP was used for the dequenching of Calcein-bound iron. Quantification of FeRhoNox-1 and Calcein-AM mean fluorescence intensity (MFI in arbitrary units (a.u.)) by FACS analysis are shown. n = 4-6. **c,** Comparison of IPA upstream regulator analysis of DEG in Fbxl5-KO HSC (GSE93649), previously defined mouse HSC ageing signatures (GSE166674), and ageing-associated proteins in human plasma (PMID33089916). Z-scores of top 30 upstream regulators are shown. **d,e,** Quantification of IRE-containing transcripts of murine HSC ageing signatures (**d**), or DEG of human aged (65 to 75 year old) compared to young (18 to 30 year old) HSC (**e,** GSE10440). Significance of enrichment was calculated by hypergeometric test. **f,** Comparison of the expression of IC-treated HSC versus control HSC in scRNA sequencing, with up- or down-regulated genes in murine aged HSC signatures. Signature scores were calculated with *Seurat* package. **g,** *GSEAPreranked* analyses of IC-treated HSC scRNA-seq data using gene sets of up-regulated signatures in murine aged HSC (**g**). **h,** *GSEAPreranked* analyses of IC-treated HSC scRNA-seq data using gene sets of ‘*MEGAKARYOCYTE_DIFFERENTIATION*’ retrieved from the gene ontology (GO) database, as well as significantly up-regulated genes in aged versus young HSC that are positive for von Willebrand Factor (vWF+). Single cell RNA sequencing data of aged and young HSC were obtained from GSE70657; HSC with vWF expression > 3 were defined as vWF+. **i,** Expression of genes in leading edge of GSEA in (**g**) was compared in aged HSC, IC-treated HSC, as well as Tip60 KO LSK or Myc dKO HSC. Genes implicated in platelet activation or production, as well as hematopoietic differentiation are indicated. **j,** Scheme of transplantation assay to evaluate the effect of iron chelation on aged HSC. HSC were isolated from aged mice following 2-week treatment of iron chelator (DFO) or vehicle control (PBS), and transplanted into congenic recipients (500 HSC each) via retro-orbital injection. Donor cell chimerism was assessed by FACS analysis 16 weeks after transplantation. **k,l,** Donor cell (CDD45.2^+^) chimerism (**k**) and frequency of donor-derived HSC (**l**) in recipient bone marrow 16 weeks after transplantation. **m,n,** Frequency of CD41^+^ HSC (**m**), and megakaryocyte-primed progenitors (CD3^-^/CD4^-^/CD8^-^/ B220^-^/CD11b^-^/CD41^+^; **n)** within the donor-derived cells. **o,p,** Quantification of megakaryocytic differentiation of aged HSC upon IC (IC1: DFO) exposure alone or in combination with Tip60 or Myc inhibition for 7 days. Representative morphologies of megakaryocytic colony-forming unit (CFU-Mk) are shown (**o**). Quantification of CFU-Mk colonies across different conditions were normalized to 2000 HSC plated (**p**). n = 8; 3 independent experiments. If not specified otherwise, data are mean ± SEM (**a, b, k-n,** and **p**). Significance p-values, indicated as *p < 0.05, **p < 0.01, ***p < 0.001, ****p < 0.0001, were calculated using Student’s t-test (unpaired: **a, b, f**, and **k-n**; paired: **p**).

We next determined whether IC exposure increased HSPC due to altered intracellular iron availability or as a consequence of iron regulatory pathway activation. Pharmacological inhibition of the limited iron response by employing a CD71-blocking antibody along with a highly specific inhibitor for lysosomal degradation of ferritin (Vps34i) (**Extended Data Fig. 4i**) blocked the HSPC stimulatory effects of IC treatment (**Fig. 1l**), as did heterozygous ablation of *Fth1* (**Extended Data Fig. 4j,k**). This set of data demonstrates that the increase in HSPC following IC-treatment is attributable to the activation of the limited iron response which promptly replenishes the intracellular LIP upon iron chelation.

## Fatty acid oxidation drives HSC expansion upon limited iron response activation

To better understand the precise molecular mechanism leading to increased HSC numbers after iron homeostasis activation, we performed a comprehensive molecular analysis of primary HSC and HPC-7 cells following IC exposure. Compared to vehicle treated cells, IC exposure led to transcriptomic changes in HSC consistent with metabolic reprograming - affecting glycolysis as well as lipid, particularly arachidonic acid-driven, metabolism as early as 12 hrs and lasting for at least 48 hrs post treatment (**Fig. 2a, Extended Data Fig. 5a-f, Extended Data Tables 2-5**); we found alterations in steroid hormone receptor (peroxisome proliferator activated receptor (PPAR alpha) / retinoid X receptor (RXR alpha), Glutathione redox reactions and phosphatidylglycerol biosynthesis pathways (**Extended Data Fig. 5e, Extended Data Table 4**), associated with lipid metabolism and transport, amino acid metabolism and regulation of prostaglandin concentration (**Extended Data Fig. 5f, Extended Data Tables 4,5**). Metabolomic analysis of IC-treated HPC-7 cells as well as primary HSC-enriched cell populations confirmed these observations, and further suggested that the increase in fatty acid mobilization, lipolysis and fatty acid oxidation was at least in part dependent on ferritinophagy (**Extended Data Fig. 5g-j, Extended Data Tables 6-9**). Direct quantification of fatty acid oxidation (FAO) showed increased abundance of a key FAO enzyme, Hadha, in primary HSC and other immature HPSC populations (**Fig. 2b**), as well as in HPC-7 cells (**Extended Data Fig. 5k-m**) upon IC exposure compared to mock controls; we also found elevated basal and maximal FAO rates by Seahorse™ analysis in hematopoietic Ba/F3 cells upon IC exposure of compared to vehicle controls, which was reversible upon lipolysis blockage (**Fig. 2c, Extended Data Fig. 5n,o**). IC exposure did not trigger detectable alterations in the overall ATP production while we noticed a moderate, albeit statistically non-significant increase in FAO-driven ATP generation (**Extended Data Fig. 5p**).

We tested whether increased FAO contributed to the stem cell stimulatory effects of IC treatment. Pharmacological inhibition of mitochondrial FAO using carnitine palmitoyltransferase-1 (CPT-1) inhibitor, etomoxir, prevented HSC expansion upon IC exposure (**Fig. 2d, Extended Data Fig. 5q**). We also detected an increase in asymmetric cell divisions driving IC-mediated HSC pool expansion (**Fig. 2e**), in line with prior work demonstrating increased asymmetric cell divisions upon FAO stimulation in HSC^50^. Together, these data show that activation of the limited iron response triggers metabolic reprogramming, increasing lipid mobilization and fatty acid oxidation which expands the HSC population.

## Arachidonic acid driven FAO drives iron dependent HSC expansion

We next investigated which fatty acid pool served as a source for the observed increase in mitochondrial β-oxidation. We detected a transient increase in Cyclooxygenase 1 (COX-1 or prostaglandin G/H synthase 1/2 (Ptgs1)) in HSC 8 to 16 hrs upon IC treatment compared to vehicle controls (**Extended Data Fig. 6a**), as well as elevated intracellular levels of the polyunsaturated fatty acid (PUFA) arachidonic acid (AA) (**Fig. 2f**). Consistently, our molecular profiling showed alterations associated with increased AA release, prostaglandin and eicosanoid production, as well as catabolism of AA (**Fig. 2a, Extended Data Fig. 5h,i**). Analysis of proteome profiling data^62^ demonstrated the robust to high expression of several AA metabolic pathway enzymes in HSC; notably, enzymes facilitating intracellular AA mobilization (Pla2g4a) and AA-dependent synthesis of eicosanoids (Cyclooxygenases 1 and 2 (COX-1/2 or prostaglandin G/H synthase 1/2 (Ptgs1/2), Thromboxane A synthase 1 (platelet, cytochrome P450, family 5, subfamily A; Tbxas1))^63^ show particularly high protein levels in steady state HSC (**Extended Data Fig. 6b**), which is in line with the demonstrated regulatory function of AA derived lipid metabolites and reactive oxygen species (ROS) formation in HSC self-renewal, homing and engraftment^64–66^.

We also detected a prompt, yet transient increase in the abundance of Long-chain fatty acid-CoA ligase 4 (Acsl4) upon IC exposure, which appeared dose dependent (**Fig. 2g**). Acsl4 is an essential acyl-CoA synthetase catalysing the conversion of long-chain fatty acids, preferentially AA, into their active form, acyl-CoA, for lipid synthesis and degradation via β-oxidation^67^. We predicted that increased Acsl4-mediated activation of AA contributed to the elevated rates of FAO driving HSC expansion upon intracellular iron chelation. RNAi mediated knock-down of *Acsl4* (**Extended Data Fig. 6c,d**) reduced basal FAO at steady state, as well as basal and maximal FAO upon IC treatment compared to non-targeting controls (**Fig. 2h, Extended Data Fig. 6e,f**) in BaF/3 cells; importantly, *Acsl4* ablation also prevented IC-driven primary HSPC expansion (**Fig. 2i**). This set of results strongly supports a role for AA-driven β-oxidation in the IC-mediated increase of functional stem cells.

## Tip60 controls iron dependent gene expression programs in HSC

Lipid-derived acetyl-CoA can serve as a major carbon source for histone acetylation, especially when glucose and glutamine metabolism are impaired.^68^ Whether and how lipid carbon contributes to gene regulation in HSC has been unknown. Given the observed increase in FAO and concomitant decrease in glucose pathways following IC exposure (**Fig. 2a-c, Extended Data Fig. 5a,c**)^44^, we predicted that increased FAO would lead to functionally relevant changes in acetyl-CoA production and protein acetylation in HSC. In support, we found that addition of a pharmacologic inhibitor of ATP citrate lyase (ACLY), a transferase that catalyses the conversion of citrate to acetyl-CoA in the cytoplasm and nucleus of cells,^69^ partially reversed the stem cell stimulatory effects of IC treatment (**Fig. 3a, Extended Data Fig. 7a**).

We furthermore found evidence of increased histone acetyltransferase activity, specifically MYST family member Tip60 (KAT5) - the acetyl-CoA N-acetyltransferase and catalytic subunit of the NuA4 complex upon IC treatment (**Fig. 3b, Extended Data Fig. 7b,c**). Tip60/NuA4 activates selected genes through acetylation of core histone (H) 4 at lysines (K) 5, 8 and 12, H2A at K5,^70, 71^ and histone variants H2AX and H2AZ^72^. Tip60/KAT5 is an essential regulator of cellular homeostasis and stress response, and has been implicated in the maintenance and renewal of embryonic stem cells^73^ and HSC^72^. Using gene set enrichment analysis, we found a significant positive enrichment of H2AZac, H4K5ac, H4K8ac, H4K12ac, H4K16ac regulated genes at a genome wide level (**Fig. 3c, Extended Data Fig. 7d**). Interestingly, we also noted a strong enrichment of genes harbouring transcription activating mono- and tri methylation marks on H3K4^74^, as well as transcriptional fidelity-enhancing trimethylated H3K36^75^ in HSC following IC treatment which appeared to be restricted to Tip60 occupied genes (**Fig. 3c**). These observations strongly suggested that IC treatment increases histone acetylation and enhances Tip60-mediated gene regulation in HSC.

Gene-specificity of the Tip60/NuA4 complex is conferred by several transcription factors, including Myc^76^ which is known to interact with the complex to drive H3 and 4 acetylation in mouse embryonic stem cells^77^, and H2AZ acetylation in LSK-HSC^72^. While we did not detect significant upregulation of *c-Myc* in the majority of stem cells upon IC exposure (**Extended Data Fig. 7e**), we found enrichment of Myc-regulated genes, including known Tip60 targets differentially expressed (**Extended Data Fig. 7b**) and consistent with an increase in Myc activity in HSC following IC treatment (**Extended Data Fig. 7f-h**). We furthermore uncovered enrichment of c-Myc regulated targets within the subset of direct Tip60 genes differentially expressed upon IC treatment in HSC (**Fig. 3d, Extended Data Fig. 7i**); analysis of transcriptional changes in HSC isolated from a mouse model of iron overload (induced by *Fbxl5* deletion)^43^ further showed a large overlap of pathways shared with differential gene expression found in *Myc* or *Tip60*-deficient HSC (**Extended Data Fig. 7j,k**). This suggested that Tip60 and Myc cooperate in the regulation of gene expression following IC treatment of HSC. To test this, we evaluated whether inhibition of Tip60 and Myc would curtail the HSC stimulatory effects of IC. Fluidigm™-based gene expression assessment of a set of predicted Tip60/Myc regulated genes showed attenuation of several gene expression alterations upon IC exposure in HSC co-treated with highly selective pharmacologic inhibitors; these included Mevalonate kinase (*Mvk*) and Cyclin-dependent kinase 4 inhibitor D (*Cdkn2d*) (**Fig. 3e**), two targets of the Mediator co-activator complex component CDK19 (Cyclin Dependent Kinase 19), a negative regulator of lipogenesis and CD71 endocytosis^78, 79^, and whose targets were enriched in HSC-specific Tip60 regulated gene sets (**Fig. 3d, Extended Data Fig. 7l**). Moreover, we confirmed upregulation of High Mobility Group AT-Hook 1 (*Hmga1*) (**Fig. 3e**), a chromatin modulator well expressed in HSC^80^, which is known to cooperate with the Mediator complex to drive gene-specific regulation^81^, including the suppression of Numb^82^ which induces differentiation in HSC^25, 83^. At the cell functional level, pharmacological inhibition of Tip60 or Myc showed a complete inhibition of the stem cell stimulatory effects of IC while exerting no effect on steady state HSC (**Fig. 3f,g, Extended Data Fig. 7m**). Together, these data show that iron limitation induces a gene-specific increase in Tip60-mediated histone acetylation and enhanced gene expression control orchestrated by Myc and likely CDK19/Mediator (**Extended Data Fig. 7n,o**) which reinforces hematopoietic stem cell identity.

## Cytoplasmic iron loading in ageing HSC

Loss of iron homoeostasis occurs during ageing and manifests in (1) iron deficiency anaemia - originating from low circulating body iron due to insufficient iron consumption or chronic inflammation^84^; (2) perturbed intracellular iron partition and mitochondrial function caused by decreased iron sulphur cluster and heme biosynthesis^85–89^, as well as (3) intracellular iron loading in several tissues^90–94^. We next assessed how ageing-associated loss of iron homeostasis may affect tissue-specific stem cells.

Quantification of the cytoplasmic LIP showed elevated levels of chelatable Fe^2+^ as well as Fe^3+^ pools in purified HSC from aged (22 to 24 mos.) vs. control cells from young (2 to 3 mos.) mice (**Fig. 4a,b**); a similar expansion of the LIP was detected in aged multipotent and committed progenitor cells compared to young controls (**Fig. 4a,b, Extended Data Fig. 8a,b**). Assessment of peripheral blood iron parameters showed no signs of an overt systemic iron overload (**Extended Data Fig. 8c,d**). We next assessed whether an elevated LIP contributed to ageing-associated HSC dysfunction.

Comparative analysis of differential gene expression in aged mouse HSC and plasma protein levels in elderly humans (vs. young controls) not only uncovered a substantial overlap of dysregulated molecular pathways^95, 96^; it also showed strong similarly with molecular alterations found in *Fbxl5* deficient HSC^43^ (**Fig. 4c**). Furthermore, IRE-motif analysis revealed a highly significant enrichment of IRE-motif containing transcripts found aberrantly expressed in aged (vs. young) mouse as well as human HSC^97^ (**Fig. 4d,e, Extended Data Fig. 8e**). We also noted that ageing associated gene expression alterations in HSC were inversely correlated with those found upon IC treatment (**Fig. 4f,g, Extended Data Fig. 8f-h**). These changes particularly affected genes with roles in megakaryopoiesis, and were also found enriched in mostly dormant megakaryocytic-biased, van Willebrandt factor positive (vWF+) HSC (Mk-HSC; known to expand during ageing^98, 99^) (**Fig. 4h, Extended Data Fig. 8i**), which included several Tip60/Myc regulated genes (**Fig. 4i, Extended Data Fig. 8j**). This suggested that IC treatment may reverse at least some of the ageing-associated phenotypes in HSC.

## Iron chelation restores function of aged HSC

We next subjected old mice to a 14-day IC (or mock control) treatment regimen and, upon its completion, competitively transplanted equal numbers of highly purified phenotypical HSC into lethally irradiated young congenic recipients (**Fig. 4j**). Recipients of IC-treated donor stem cells showed an increase in overall donor cell chimerism (**Fig. 4k**) and an increased number of donor-derived phenotypical HSC (**Fig. 4l**). Notably, we found a concomitant decrease in phenotypic Mk-HSC with a known ageing-associated aberration – cell surface presentation of glycoprotein (Gp) IIb/IIIa integrin (CD41) (**Fig. 4m**), a receptor for fibrinogen which is typically expressed on megakaryocytic (Mk) progenitors and platelets^100^. In line, recipients of IC-treated aged HSC showed lower donor-derived CD41 expressing Mk progenitor cell frequencies compared to mock treated control cell recipients (**Fig. 4n**). *Ex vivo* IC treatment followed by quantification of megakaryopoiesis commitment showed alleviation of the megakaryocytic differentiation bias of aged HSC (**Extended Fig. 8k**) upon iron limitation (**Fig. 4o,p**); notably, pharmacological inhibition of Tip60 or c-Myc partially reversed the IC-induced mitigation of aberrantly increased megakaryocytic differentiation bias of aged HSC (**Fig. 4p, Extended Data Fig. 8l-n**). Together, these observations demonstrate that IC treatment can restore deregulated iron-dependent molecular programs driving the dysfunction of aged stem cells.

## Discussion

In this study, we provide evidence supporting the pool of redox active, readily accessible labile iron as a key cellular rheostat allowing HSC to orchestrate metabolic and gene regulatory pathways that reinforce stem cell identity. We also demonstrate that cytoplasmic iron loading is a key factor in HSC dysfunction during ageing.

Intact iron homeostasis is known to ensure proper hematopoietic progenitor cell expansion and differentiation^101^, to curtail reactive oxygen species accumulation and limit oxidative damage-induced elimination of haematopoietic stem cells^43^. Here, we have delineated several new molecular principles governing iron-dependent somatic stem cell function (**Extended Data Fig. 9**): First, we uncovered that mitotic stem cells activate a limited iron response during mitosis and upon experimental acute iron limitation. This demonstrates that, like in metabolically highly active and proliferating cells^46, 102^, the size of the LIP is tightly regulated and closely linked with the functional state of quiescent and metabolically less active stem cells^26, 49–52^. It also shows that metabolic demands during mitosis provide an effective trigger for the activation of iron homeostasis pathways in stem cells. Second, our data show that the LIP serves as a molecular switch to transiently shift heavily iron-dependent energy production^103, 104^, including glycolysis (of quiescent stem cells) and oxidative phosphorylation (of proliferating HSC), to metabolic pathways requiring less iron - particularly fatty acid oxidation, which is known to expand the HSC pool^50^. Third, our data strongly suggest increased lipid carbon-fuelled histone acetylation and epigenetic regulation to follow iron homeostasis pathway activation. This finding strengthens the emerging paradigm of a tight interconnection between lipid metabolism and epigenetic control in stem cells^105, 106^; it also provides mechanistic insights into metabolism-driven epigenetic regulation of HSC identity. Fourth, we demonstrate that cytoplasmic iron loading occurs in aged stem cells, blunting iron-dependent HSC gene regulation, and driving the dysfunction of aged stem cells.

Given that many somatic stem cells share principle cellular and molecular regulatory circuits^107^, it is possible that LIP serves as a general rheostat in various adult stem cells. As loss of iron homeostasis is observed in a large fraction of the elderly^108^, in patients with chronic inflammation^109^ or cancer^110^, our findings will have implications in understanding and therapeutic mitigation of altered stem cell function in a wide range of degenerative and malignant pathologies.

## Methods

### Chemicals and Reagents

Eltrombopag (EP, pure compound provided by Novartis) was reconstituted in sterile distilled water as 1 mg/ml stock and was stored at ambient temperature, light-protected for up to 2 weeks. Deferoxamine (DFO, Sigma) was freshly prepared with sterile distilled water for every experiment. Anti-CD71 blocking antibody (MCA2396EL, Bio-Rad) and Etomoxir (Sigma) were stored at −20°C in aliquots of 1 mg/ml and 10 mM stocks respectively. Inhibitors for VPS34 (RGNCY-0041/0042, Reagency), Myc (10058-F4, Selleck) and Tip60 (TH1834, Axon) were reconstituted in dimethyl sulfoxide (DMSO) and stored at −80°C until use. Iron chemo-sensors Calcein-AM (Invitrogen) and FeRhoNox-1 (Goryo Chemical) were stored light-protected at −20°C and were freshly reconstituted in DMSO for every experiment. ATP citrate lyase (ACLY) inhibitor SB 204990 (Tocris, Minneapolis, MN) and Senexin A (Selleck Chemicals, Houston, TX) were reconstituted in DMSO with a stock concentration of 10 mM and stored at −20°C until use. Cortistatin A was obtained from the laboratory of Matthew Shair at Harvard University, and reconstituted in DMSO with a stock concentration of 1 mM and stored at −20°C until use.

### Mice and cell lines

C57/BL6 (stock number: 000664), Pepc/BoyJ mice (002014), and *Fth1^LoxP^* mice (018063) were purchased from Jackson Laboratories and housed in animal facilities at the Albert Einstein College of Medicine. Male and female mice at the age of 6-10 weeks were utilized for the experiments. All experiments were approved by the Institutional Animal Care and Use Committee of the Albert Einstein College of Medicine (Protocol# 0000-1015). All procedures were performed in accordance with guidelines from the Institutional Animal Care and Use Committee of the Albert Einstein College of Medicine. The murine multipotent progenitor cell line HPC-7 was provided by Dr. Omar Abdel-Wahab (Memorial Sloan Kettering Cancer Center). Murine BA/F3 and human 293T cell lines were purchased from ATCC. All cell lines were routinely monitored for and testing negative for *mycoplasma*.

### Cell culture

#### Cell lines

HPC-7 cells were passaged in IMDM with 5% fetal bovine serum (FBS), 1% penicillin/streptomycin, 1% sodium pyruvate, 6.9 ng/mL monothioglycerol (Sigma) and 100 ng/ml recombinant mouse (rm) SCF^111^. BA/F3 cells were cultured in RPMI 1640 with 10% FBS, 2 ng/ml rmIL-3 and 1% penicillin/streptomycin^112^. 293T cells were cultured in DMEM with 10% FBS and 1% Penicillin/Streptomycin. **Primary cells.** Primary HSC were isolated by cell sorting on a Moflo Astrios EQ (Beckman Coulter). Lin^−^Sca-1^+^c-Kit^+^CD150^+^CD48^−^ (HSC) or Lin^−^Sca-1^+^c-Kit^+^CD150^+^CD48^−^CD34^−^ (CD34^−^HSC) populations were cultured in Myelocult M5300 (STEMCELL Technologies) with 100 µg/ml Primocin (Invivogen) supplemented with 100 ng/ml recombinant mouse (rm) SCF (R&D Systems) and 50 ng/ml rmTPO (R&D Systems). For lineage-negative (Lin^−^) and Lin^−^c-kit^+^ (LK) cells, 50 ng/ml of rmSCF and rmTPO were used, with 20 ng/ml rmIL-3 (R&D Systems), 50 ng/ml Flt3-Ligand (R&D Systems) supplemented to the culture. Cells were maintained at 37 °C and 5% CO_2_ unless otherwise specified.

### Flow cytometry analysis and sorting

#### Cell preparation and sorting

Isolation of mononuclear cells (MNC) from mouse bone marrow was performed as previously described^44^. Briefly, MNC were lineage-depleted using 1:200 dilution of anti-mouse CD4, CD8, B220, CD19, Ter119 and Gr-1, all biotin-conjugated, rotating at 4°C for 30 min. Cells were washed and then stained with triple-washed anti-IgG magnetic beads (Untouched Mouse T Cells Kit, Thermo Fisher) rotating at 4°C for 30 min. Cells were washed and then depleted of lineage-positive cells by passing through a magnetic separation column (MACS LD Column, Miltenyi Biotec) loaded on a DynaMag-5 Magnet (Invitrogen). Lineage-negative cells were then stained for 30 min on ice with stem and progenitor cell markers (Sca-1, c-Kit, CD150, CD48, CD34; 1:100). A detailed description of mouse antibodies can be found in **Extended Data Table 10**. Lin^−^Sca-1^+^c-Kit^+^CD150^+^CD48^−^ (HSC) or Lin^−^Sca-1^+^c-Kit^+^CD150^+^CD48^−^CD34^−^ (CD34^−^HSC) and Lin^−^Sca-1^+^c-Kit^+^ (LSK) cells were sorted on MoFlo Astrios EQ (Beckman Coulter). **Quantification of CD71.** Lin-cells were stained for cell surface markers defining stem and progenitor cells (HSPC), as well as cell surface transferrin receptor (CD71, eBioscience). Stained cells were then fixed with Cytofix/Cytoperm buffer (BD Bioscience, 554722) for 20 min on ice, washed twice with Perm/Wash buffer (BD Bioscience, 554723), re-suspended in Perm/Wash buffer containing anti-CD71 antibody (abcam, ab84036) and incubated for 1 hour at room temperature (RT). After 2 washes with Perm/Wash buffer, cells were re-suspended in Perm/Wash buffer containing Alexa Fluor 647 goat anti-rabbit IgG (H+L) (Invitrogen, A21245) for 30 min at RT. Cells were washed twice with Perm/Wash buffer and subjected to FACS analysis. **Quantification of Hadha**. For primary cells, lineage-depleted MNC were treated with iron chelators (10 µM DFO) for 24 hours, followed by staining with FACS antibodies against surface markers for HSPC. Thereafter, stained cells were fixed and permeabilized with Cytofix/Cytoperm buffer for 20 min on ice. For HPC-7, cells were treated with iron chelators (10 µM DFO or 5 µg/ml EP) for 24 hours before fixation and permeabilization with Cytofix/Cytoperm buffer. Intracellular staining with anti-Hadha antibody (abcam, ab203114) for 1 hour at RT, followed by secondary staining with Alexa Fluor 647 goat anti-rabbit IgG (H+L) (Invitrogen, A21245) for 30 min at RT. **Calcein-AM**. Intracellular iron levels were measured by calcein-acetoxymethyl ester (calcein-AM, Invitrogen) assay as previously described^44, 113^. Primary bone marrow MNC or HPC7 cells were preloaded with 50 nM calcein-AM (Invitrogen) at 37°C for 5 min, before the treatment with vehicle control or iron chelators for 2 to 6 hours at 37°C. Treatments were performed in complete media (**Extended Data Fig. 3h**, 4 to 6 hr exposure) or PBS supplemented with 1% polyvinyl alcohol (**Extended Data Fig. 8b**, 2 hr exposure) to model the ability of cells to access transferrin-bound iron extracellularly. For quantifying Fe^3+^ cytoplasmic pool, a ferric iron-specific chelator (eltrombopag, Novartis) was used to liberate calcein-bound Fe^3+^ iron (**Extended Data Fig. 8b**). For primary MNC, cells were stained with antibodies against cell surface markers for HSPC on ice for 15 min. Thereafter, intracellular calcein fluorescence was analysed by flow cytometry. The relative size of the labile iron pool (LIP) was calculated as the difference in mean fluorescence intensity (ΔMFI) between the vehicle and iron chelator-treated cells^114^. **FeRhoNox-1**. Intracellular iron levels were also measured with FeRhoNox-1 fluorescent imaging probe specific for ferrous iron (Goryo Chemical). For primary cells, lineage-depleted MNC were incubated with 20 μM FeRhoNox-1 at 37°C for 1 hour. Antibody cocktail for HSPC surface staining was added in the last 10 min of FeRhoNox-1 incubation. For HPC-7, cells were pre-treated with VPS34 inhibitor (Reagency, RGNCY-0042) for 2 hours before exposure to vehicle or iron chelator (10 μg/ml eltrombopag) for 1 hour at 37°C. Thereafter, treated cells were washed with PBS, and incubated with 20 μM FeRhoNox-1 at 37°C for 1 hour. Cells were washed twice with PBS and subjected to FACS analysis. Unless otherwise specified, fluorescent signals were acquired with BD FACS Aria II system (Becton Dickinson) and analysed with FlowJo V10.2.

### Single molecule RNA FISH

To design *Tfrc* mRNA-specific probes for sequential single molecule FISH (smFISH), full length transcript of *Tfrc* (NM_011638) was used as input for PaintSHOP^115^ to retrieve 22 primary targeting sequences (30-40bp, **Extended Data Table 11**), separated by at least 10bp. Putative sequences were then screened for off-target activity using NCBI Blast (https://blast.ncbi.nlm.nih.gov/Blast.cgi) against mouse transcriptome. Selected sequences were then concatenated on the 5′ and 3′ end with flanking readout 20mer sequences (GTTTGAAGATTCGACCTGGA), generating a final ‘primary probe’.

SmFISH immunofluorescence staining procedure and analysis were performed as described previously^58, 116^. Briefly, treated and control cells were attached to coverslips using biotinylated anti-CD44 coating^117^. Residual media was washed with PBS, and cells were fixed in 3.2% PFA (Electron Microscopy Sciences), diluted in PBS with 1 mM MgCl_2_ (PBSM), for 10 minutes at room temperature. Cells were then permeabilized in 0.1% Triton X-100 in PBSM for 10 minutes. After washing with PBSM, cells were incubated at room temperature with 30% prehybridization buffer (30% formamide, 2X saline-sodium citrate buffer) for 30 minutes. Primary hybridization was done in 30% hybridization buffer consisting of 10% dextran sulfate, 30% formamide, 2X saline-sodium citrate (SSC), 2 mM VRC, 10 μg/ml sheared ssDNA from salmon sperm, 10 μg/ml E. coli tRNA, 10 μg/ml molecular grade bovine serum albumin (BSA), and 200 ng of primary probe mixes, overnight at 37 °C. Thereafter, cells were washed twice with 30% pre-hybridization buffer for 20 min at 37 °C and once with 2X SSC. Cells were then post fixed in 3.2% PFA in PBSM for 10 min, followed by washing in 2X SSC. Primary stained cells were incubated with 10% prehybridization buffer (10% formamide, 2X SSC) for 10 min at 37 °C and stained with 10% dextran sulfate, 10% formamide, 2X SSC, 2 mM VRC, 10 μg/ml sheared ssDNA from salmon sperm, 10 μg/ml E. coli tRNA, 10 μg/ml molecular BSA, and 10 ng Cy5-labelled readout probe of 20mer readout probe (RO2-Cy5) for *Tfrc* gene for 3 hrs at 37 °C. Cells were then washed twice for 10 min in 10% prehybridization buffer, followed by a final wash in 2X SSC. Before the immunostaining for pS10H3, cells were again fixed with 3.2% PFA in PBS for 10 min at room temperature (RT). Cells were washed once with PBS for 5 min at room temperature. Blocking was performed in blocking buffer (PBS, 1% RNAse-free BSA, 0.2% Triton X-100) for 30 min at room temperature. Next, cells were incubated with the primary antibody (1:200 mouse anti-pS10H3, Cell Signaling Technology) in antibody dilution buffer (PBS, 0.1% BSA, 0.1% Triton X-100) overnight at 4 °C. Next day, cells were washed thrice with PBS for 5 min at RT before incubation with the secondary antibody (1:200 rabbit anti-mouse AlexaFluor 488, Cell Signaling Technology) in antibody dilution buffer for 1 hour at RT. Excess antibody was removed by washing cells with 1xPBS for 5 min at RT. Cells were then mounted in Prolong Diamond Antifade reagent plus DAPI (Invitrogen). Images were acquired using oil immersion 100X objective on an epifluorescence Olympus Digital Station 6 microscope. Exposure times were 1000 ms, 50 ms, 100 ms for Cy5, AlexaFluor 488 and DAPI respectively. Z stacks spanning the entire volume of the cells were acquired by imaging every 300 nm along the z-axis. Acquisition control of the microscope was achieved using IPLab software. For data analysis, single molecule mRNA and transcription site detection was performed using freely available and MATLAB-written software FISHquant, by 3D Gaussian fitting of thresholded spots, implemented in MATLAB R2020b^117^. Further experimental details, validation, and discussion of this methodology in the hematopoietic system can be found in ^58^.

### Stroma-free long-term culture-initiating cell assay

Conditioned media derived from mouse stromal cells were collected as previously described^44^. Long-term culture-initiating cell (LTC-IC) assays were performed as previously described^44^ for the assessment of the frequency of functional stem cells *ex vivo*. Briefly, limiting dilutions of LSK cells were FACS-sorted into 96-well plates containing Myelocult M5300 and conditioned media at 1:1 ratio, supplemented with 50 ng/ml rmSCF, 50 ng/ml rmTPO, 1 μM hydrocortisone (STEMCELL Technologies), and 200 μg/ml Primocin. After 4 weeks of culture at 32°C, 5% CO_2_, limiting dilutions of LSK cells and their respective replicate wells were subjected to methylcellulose colony assay with HSC007 (R&D Systems) at 37°C, 5% CO_2_ for 1 week. Colony forming units were identified and scored using inverted light microscope. Stem cell frequency was estimated using extreme limiting dilution analysis (ELDA) algorithm^118^.

For inhibiting the activation of iron homeostasis regulatory pathways upon intracellular iron reduction, a CD71-blocking antibody (Bio-Rad, MCA2396EL)^119^ and VPS34 inhibitor (Reagency, RGNCY-0041)^120^ were used to simultaneously block iron uptake and mobilization, respectively. For inhibiting fatty acid oxidation, etomoxir (Sigma, E1905) was used to irreversibly block mitochondrial carnitine palmitoyltransferase-1^121, 122^. For inhibiting Myc, 10058-F4^123^ (Selleck, S7153) was used to block the dimerization of Myc and Max. For inhibiting histone acetyltransferase activity of Tip60, TH1834 (Axon, 2339)^124^ was used.

### Colony-forming unit assays of megakaryocyte progenitors

For the detection of colony-forming unit (CFU) of mouse megakaryocyte progenitors, 2000 sorted HSC (Lin^-^/Sca1^+^/cKit^+^/CD150^+^/CD48^-^) were plated in a collagen-based medium in double chamber culture slides and cultured for 7 days at 37°C, 5% CO_2_ (MegaCult®-C Medium without Cytokines, STEMCELL Technologies). Cultures were supplemented with human recombinant 50 ng/ml TPO (Peprotech), 20 ng/ml human recombinant IL-6 (STEMCELL Technologies) and 10 ng/ml mouse recombinant IL-3 (R&D) according to manufacturer’s instructions. Staining for acetylcholinesterase content and scoring of CFU-Mk colonies were performed according to manufacturer’s protocol, and colonies with at least eight-cell cluster were scored using Inverted Infinity and Phase Contrast Microscope (Fisher Scientific). Where indicated, media were supplemented with: 10 µM DFO (Sigma), 3 µg/ml eltrombopag (Novartis), 10 µM Tip60 histone acetyltransferase inhibitor (TH1834, Axon), 25 µM c-Myc inhibitor (10058-F4, Selleck Chemicals).

### Bone marrow transplantation

Aged mice (22-24 mos) obtained from National Institutes for Aging were subjected to *in vivo* treatment of iron chelator (Deferoxamine, 50 mg/kg) or vehicle control (sterile PBS) daily for 14 days. Treatments were administered by intraperitoneal injection. Thereafter, HSC (Lin^-^/Sca1^+^/cKit^+^/CD150^+^/CD48^-^) were prospectively FACS-sorted from treated mice and pooled according to experimental group, where 500 HSC along with 1×10^6^ Sca-1-depleted CD45.2 Pepc/BoyJ BMMNC were transplanted into CD45.2 recipient mouse irradiated with 2 rounds of 500 rads irradiation. Detailed procedures for HSC transplantation were described previously^44^. Chimerism of donor-derived hematopoietic stem and progenitor, as well as lineage reconstitution were assessed at 16-weeks post transplantation and analyzed with BD FACS Aria II system (Becton Dickinson) and FlowJo V10.2. Detailed information of the antibodies used for donor chimerism analysis can be found in **Extended Data Table 10**.

### RNA extraction, quantitative PCR and microarray

RNA extraction, reverse transcription, and quantitative PCR (qPCR) were performed as previously described^44^. Briefly, RNA was extracted with the Qiagen RNeasy Micro Kit, and reverse-transcription of extracted RNA was performed using Superscript II reverse transcriptase (Invitrogen). For qPCR, 10 μl reaction volume containing 2 μl cDNA (5 ng/μl), 0.5 μl of each forward and reverse target primers (5 μM), 5μl of Power SYBR Green mix (Applied Biosystems), and 2 μl of nuclease-free water were used. Triplicate samples and five serial dilutions of standards were prepared for each target gene. Thereafter, qPCR was performed using ViiA 7 Real-Time PCR system (Applied Biosystems) according to the manufacturer’s instructions. Gene expression levels were calculated based on the standard curve with subsequent normalization to internal control (*GAPDH)*. A list of qPCR primers for target genes can be found in **Extended Data Table 11**.

For microarray, 5 to 20 ng of RNA were amplified and labelled with the GeneChip 3’ IVT Pico Kit (Affymetrix) according to manufacturer’s protocols. Labelled RNA was then hybridized to Mouse Clariom S microarrays (Affymetrix), scanned and analysed with GeneChip Scanner 3000 7G system (Affymetrix) according to standard protocols. Resulting CEL files have been deposited in the GEO database (GSE157817).

### Analysis of microarray data

Microarray data was analysed as previously described^44^. Briefly, quality assessment of microarray results was performed with Transcriptome Analysis Console 3.0 (Affymetrix). After data normalization with RMA algorithm of Oligo Bioconductor package^125^, paired, linear modelling of *limma* package was used for differential expression analysis^126^. Microarray expression probes were then annotated to gene names with annotation files obtained from the Affymetrix website (http://www.affymetrix.com).

Differentially expressed genes (DEG; fold change > 1.6 and *P* < 0.05) were subjected to Ingenuity Pathway Analysis (IPA, Qiagen) for the analysis of biological function and upstream regulators. DEG were also examined for the presence of Iron Response Element (IRE) using SIREs Web server 2.0 as previously described^127–129^. Significance of enrichment of IRE-containing genes was calculated by hypergeometric test with built-in *phyper* function of R (https://www.R-project.org).

### Single cell RNA sequencing (scRNA-seq)

FACS-sorted HSC (Lin^−^Sca-1^+^c-Kit^+^CD150^+^CD48^−^CD34^−^) were exposed to iron chelator (10µM DFO) or vehicle (H_2_O) for 48 hours. Treated cells were collected; viability of >95% was confirmed by trypan blue exclusion. 20,000 cells from each treatment group were subjected to 10x Genomics platform for library preparation using Chromium Single Cell 3′ Reagent Kits (v3) following the manufacturer’s sample preparation guide (PN CG00054 Rev B; Fluidigm). Following quality control assessment with Agilent 2100 Bioanalyzer, approximately 50 ng libraries were sequenced on the BGISEQ-500 platform with 28+8+91bp reads (read 1: 28bp; read 2: 91bp plus 8bp for index) using two lanes per sample. Sequencing data were deposited on GEO database (GSE157821).

For bioinformatic analysis, cell barcode processing, transcriptome alignment (mm10), and gene UMI counting were performed for each of the samples using the *count* module of Cell Ranger v3.0.2^129^. Samples were aggregated using the *aggr* module with default parameters. Cells with UMI counts <2,000 or >10,000 were excluded from further analysis to rule out contamination from potentially dead or non-single cells. Cells with mitochondrial mRNAs >15% were also excluded. Moreover, genes that were expressed in less than 50 cells were excluded from the expression matrix for downstream analysis. The expression data was then normalized by the total number of unique reads, multiplied by a scale factor of 10,000 and log-transformed. For phenotypic analysis, HSC were defined based on the expression of HSC markers *Hlf* and *Ly6a*^130, 131^. Cells with expression of *Hlf* and *Ly6a* higher than the 25% percentile of their average expression in all cells in both conditions (vehicle vs. iron chelator) were defined as HSC (green), otherwise non-HSC (orange). Frequencies of phenotypic HSC and non-HSC were visualized separately using Loupe browser v3.1.1 with t-SNE maps. Signature score in each single HSC was calculated with *AddModuleScore* module of *Seurat* package.

### Gene expression analysis by Fluidigm

Sorted HSC (Lin^−^Sca-1^+^c-Kit^+^CD150^+^CD48^−^) were treated with vehicle control, iron chelator (IC, 10 µM DFO), as well as IC along with inhibition of c-Myc (Myci, 50 µM 10058-F4) or Tip60 (Tip60i, 20 µM TH1834). Forty-eight hours post treatment, RNA extraction and reverse transcription were performed as described above. Taqman assays (Applied Biosystems) of target genes for Fluidigm gene expression analysis are listed in **Extended Data Table 11**. For pre-amplification of cDNA, reaction mixtures containing 1 µL PreAmp Master (Fluidigm, CA), 1.25 µL pooled 0.2x TaqMan assays, and 2.75 µL cDNA product were used. PCR amplification was performed as follows: 95°C for 2 min; 14 cycles of 95°C for 15 sec and 60°C for 4 min. No template control (NTC) was included in the pre-amplification as negative control. For sample Pre-Mix solution preparation, 2.25 µL pre-amplification product was mixed with 2.5 µL TaqMan Fast Advanced Master Mix (Applied Biosystems) and 0.25 µL 20X GE sample loading reagent (Fluidigm). Thereafter, sample Pre-Mix solutions were transferred to a 96-well plate. For 10X TaqMan assay preparation, 5 µL of each 20X TaqMan assay was diluted with 5 µL 2X assay loading reagent (Fluidigm), followed by transferring to a 96-well plate. Thereafter, plates containing sample Pre-Mix solutions and TaqMan assays were loaded onto 96.96 Dynamic Array IFC (Fluidigm) in Biomark HD system (Fluidigm). Each of the sample Pre-Mix solutions was mixed with each TaqMan assay in the IFC by Biomark system, and qPCR reactions were performed according to manufacturer’s protocol.

Data were analysed with Fluidigm Real-Time PCR Analysis Software v4.5.1 to obtain the Ct values of genes in each sample. For differential expression analysis, Delta Ct (ΔCt) of each gene was calculated by comparing the Ct value to the internal control *Actb*. Delta delta Ct (ΔΔCt) and fold change (2^-ΔΔCt^) were calculated by comparing ΔCt in each treatment condition to ΔCt in control.

### Immunofluorescence staining

The following primary antibodies were used for immunofluorescence: anti-Ncoa4 (Santa Cruz Biotechnology, sc-15984)^59^, anti-Numb (abcam, ab4147)^132^ and anti-CD71 (abcam, ab84036)^133^. The secondary antibodies used were: Alexa Fluor 594 donkey anti-goat IgG (H+L) (Invitrogen, A-11037) and Alexa Fluor 488 goat anti-rabbit IgG (H+L) (Invitrogen, A-11008).

*Cell culture and pre-treatment*: HSC (Lin^−^Sca-1^+^c-Kit^+^CD150^+^CD48^−^) from wildtype C57BL/6 mice were sorted into 16-well chamber slides (Thermo Fisher Scientific) coated with RetroNectin (Takara Bio). HSC were cultured in M5300 media with 100 µg/ml Primocin supplemented with 100 ng/ml rmSCF and 50 ng/ml rmTPO. <u>Ncoa4-mediated</u> <u>ferritinophagy assessment</u>: HSC were subjected to either vehicle (H_2_O) or iron chelator (DFO, Sigma) treatment for 48 hours. For rescue, VPS34 inhibitor (Reagency, RGNCY-0041) was added along with iron chelator. To allow the detection of Ncoa4 foci, chloroquine (10 µM, Sigma) was added in the last 4 hour of culture to impair degradation of autophagosomes. <u>HSC asymmetric cell division quantification</u>: HSC were treated with either vehicle (H_2_O) or iron chelator (EP) for 48 hours. For rescue, HSC were preloaded with 100 µg/ml of ferric ammonium citrate (FAC) for 1 hour at 37°C prior to iron chelator treatment.

*Cell staining and imaging*: Cells were fixed with 4% PFA for 10 min and permeabilized with 0.15% Triton X-100/PBS for 5 min at RT. Cells were then blocked in 2% BSA/0.15% Triton X-100 in PBS for 1 hour at RT and incubated overnight at 4°C with the first antibody diluted in 1% BSA/0.15% Triton X-100 followed by 45 min incubation at RT with fluorescence-conjugated secondary antibodies. Cells were washed 3 times with PBS and mounted with Prolong Gold (Molecular Probes) containing 1 μg/ml DAPI. All the images were acquired with a Confocal microscope (SP5, Leica) using 63.0x 1.40 NA oil objective and the Leica LAS-AF software. Image processing and analysis were performed using ImageJ (https://imagej.nih.gov/ij/).

### Metabolomics

Metabolomic profiling and analysis were performed as previously described^44^. *Ex vivo* treated human CD34^+^ cells, as well as murine HPC-7 and LSK cells were flash-frozen in liquid nitrogen and stored at −80°C before use. Samples were either sent to the Biological Mass Spectrometry Core Facility at University of Colorado Denver, or the Einstein-Mt Sinai Diabetes Center Stable Isotope & Metabolomics Core for further processing and analysis.

Cell pellets were lysed with lysis solution (methanol:acetonitrile:water 5:3:2 v/v/v) before ice cold extraction by vortexing for 30 min at 4°C. Insoluble proteins were pelleted by centrifugation (15,000 x g for 10 min at 4°C) and supernatants were analysed using a UHPLC system (Vanquish, ThermoFisher) coupled online to a mass spectrometer (Q Exactive, ThermoFisher). Samples were resolved over a Kinetex C18 column (2.1 × 150 mm, 1.7 μm; Phenomenex) at 25°C using a 9-minute gradient at 400 μl/min from 5% to 95% B (A: water/0.1% formic acid; B: acetonitrile/0.1% formic acid). MS analysis and data analysis were performed as described before^134, 135^. Metabolite assignments were performed using a metabolomics data analyser (MAVEN). For data analysis, the level of each metabolite was normalized to internal standard (heptadecanoic acid) and cell number. Differentially altered metabolites were submitted for pathway enrichment analysis using Hypergeometric test of MetaboAnalyst 3.0^136^. Moreover, differentially altered metabolites in each treatment condition were also analysed for biological function, upstream regulators, and canonical pathways using IPA.

### Seahorse assay

Oxygen consumption rates (OCR) were measured using a 96-well Seahorse Bioanalyzer XF 96 according to the manufacturer’s instructions (Agilent Technologies). In brief, HPC-7 or *Acsl4* knockdown BA/F3 cells were cultured in their normal growth media, exposed to either vehicle or iron chelator for 16 hours. DEUP (100 μM Diethylumbelliferyl phosphate, Sigma) was used to inhibit lipolysis. To determine the fraction of oxygen consumption dependent on fatty acid β-oxidation, cells (75,000 cells/well, with 3-4 replicate wells) were plated into 180 μl KHB media (111 mM NaCl, 4.7 mM KCl, 1.25 mM CaCl2, 2 mM MgSO4, 1.2 mM NaH2PO4) and in half of the samples etomoxir (40 μM, Sigma, E1905) was added for 15-30 min before the analysis. Once in the reader, plates were sequentially injected with 1 μM oligomycin, 2 μM FCCP and 0.5 μM Rotenone. Fatty acid β-oxidation rate was calculated as the difference in oxygen consumption in the presence or absence etomoxir. Data were normalized to cell number using trypan blue exclusion and analysis was performed with Waze v2.6.0.31.

### *Acsl4* knockdown

Short hairpin RNAs (shRNAs) targeting *Acsl4* (TL502838, OriGene) were packaged in the pGFP-C-shLenti plasmid system. A nontargeting 29-mer scrambled shRNA cassette in pGFP-C-shLenti vector (TR30021, OriGene) served as a control. The sequences of *Acsl4* shRNAs can be found in **Extended Data Table 11**. For production of virus particles, lentiviral shRNA expression constructs were transfected together with packaging vectors (VSV-G, RSV-Rev and GAG-Pol) into 293T producer cells using CalPhos Transfection Kits (Takara Bio) following the manufacturer’s instructions. Supernatants were harvested after 48 and 72 hours post transfection, and concentrated by ultracentrifugation (25,000 rpm at 4°C for 2 hours). For cell lines, lentiviral transduction was performed by adding the virus supernatant to BA/F3 cells and spin-infected for 90 min at 1000 x g, 37°C. After spin-infection, the cells were exposed to the virus for 2.5 hours. Forty-eight hours post transduction, the percentage of transduced cells were analysed by FACS for GFP positivity. For primary mouse bone marrow cells, lineage-depleted MNC were enriched for c-Kit+ cells by magnetic-activated cell sorting (MACS) following manufacturer’s protocol. Briefly, Lin-cells were re-suspended in 2%FBS/PBS and incubated with FcR blocking reagent for 10 min on ice and then mouse monoclonal CD117-microbeads (Miltenyi; 130-091-224) for 15 min on ice. Prior to lentiviral transduction, Lin-c-Kit+ (LK) cells were pre-cultured for 4 hours in M5300 media supplemented with 100 μg/ml primocin, 50 ng/ml rmSCF, 50 ng/ml rmFlt3L, 50 ng/ml rmTPO, and 2 ng/ml rmIL-3. Lentiviruses for sh*Acsl4* and scrambled nontargeting control were added to RetroNectin (TaKaRa)-coated 12-well plates and centrifuged for 2 hours at 1000 x g at 32°C. After centrifugation, supernatant was discarded and pre-cultured LK cells were added immediately at a density of 500,000 cell/ml. Thereafter, the cells were spin-infected for 2 hours at 1000 x g at 37°C. After spin-infection the virus-containing media was removed and replenished with fresh M5300 media supplemented with cytokines. Forty-eight hours post transduction, GFP-positive cells were sorted on BD FACS Aria II (Becton Dickinson) and knockdown of Acsl4 was confirmed using qPCR and western blot.

### Western blot

To assess *Acsl4* knockdown, sorted primary GFP-positive cells were analysed by Western blot. For HPC-7 cell analyses, cells treated with iron chelators or vehicle control, protein expression changes were assessed for Acsl4, Cox1/Ptgs1, ferritin heavy (Fth1) and light (Flt1) chains.

Proteins were extracted from cells using RIPA buffer (50 mM Tris, pH 7.4, 1 mM EDTA, 150 mM NaCl, 1% Triton X-100, 1% deoxycholate and 0.1% SDS) supplemented with EDTA-free protease inhibitor cocktail (Sigma, 11873580001) and 1 mM PMSF. Protein concentrations were determined using Protein Assay Kit (BioRad, 5000002). Prior to loading, cell extracts were mixed with appropriate volumes of 5x protein loading buffer (10% SDS, 25% 2-Mercapoethanol, 50% Glycerol, 125 mM Tris-HCl pH 6.8, and 0.125% Bromophenol blue) and boiled at 95°C for 10 min. Subsequently, 20-30 μg of total proteins from each sample were separated by SDS/polyacrylamide gel electrophoresis in running buffer (25 mM Tris pH 8.3, 192 mM Glycine, 0.1% SDS) and transferred onto nitrocellulose (NC) membranes (Bio-Rad, 1620094) in transfer buffer (25 mM Tris pH 8.3, 192 mM Glycine) under 300 mA constant current for respective period of time depending on the molecular weight of the target proteins. NC membranes were washed once with Tris-buffered saline-Tween 20 (TBST, 20 mM Tris pH 8.3, 137 mM NaCl, 0.1% Tween 20) and blocked in 5% skim milk/TBST for 1 hour at RT. After washing three times with TBST, NC membranes were probed overnight at 4°C with the following primary antibodies: Acsl4 (Santa Cruz Biotech, sc-365230), Cox1/Ptgs1 (Cell Signaling Technologies, 4841), ferritin light chain (abcam, ab109373), ferritin heavy chain (abcam, ab65080) and actin (abcam, ab3280). Prior to and after the incubation with goat anti-rabbit (Santa Cruz Biotech, sc-2004) or goat anti-mouse (Santa Cruz Biotech, sc-2005) IgG-HRP diluted in 5% skim milk/TBST at RT for 1 hour, NC membranes were washed with TBST for three times. All blots were developed using Pierce™ ECL Western Blotting Substrate (Thermo Fisher Scientific, 32106). Signals were visualized and collected by LI-COR Odyssey Fc (LI-COR Biosciences). ImageJ (https://imagej.nih.gov/ij/) was used for quantification.

### ELISA

For the preparation of serum, peripheral blood was collected from mice into non-anticoagulant-treated tube and left undisturbed at RT to clot for 30 minutes. Clot was removed by centrifuging at 2,000 RCF for 10 minutes in a refrigerated centrifuge, and supernatant was then transferred and aliquoted into 1.5 mL clean polypropylene tubes for storage at −80°C until use. ELISA assay was performed with Mouse Transferrin ELISA kit (ab157724, Abcam, Waltham, MA) and Mouse Ferritin (FTL) ELISA kit (ab157713, Abcam) according to manufacturer’s protocols. Briefly, serum sample was diluted 1:40, or 1:100,000 with 1X diluent for the quantification of ferritin or transferrin protein, respectively. Standard samples were prepared according to manufacturer’s instructions. 100 μL diluted sample or standard control were transferred to 96 well plate strips and incubated in the dark at RT for 30 minutes. Thereafter, each well was emptied and washed four times with 200 μL 1X Wash buffer. 100 μL of 1X HRP-Antibody conjugate was then transferred to each well and incubated in the dark at RT for 30 minutes, followed by four washes with 1X Wash buffer. Thereafter, 100 μL of TMB Substrate Solution was transferred to each well and incubated in the dark at RT for 10 minutes. Reaction was stopped by adding 100 µL of Stop Solution to each well. Absorbance at 450 nM was then measured with FLUOstar Omega Microplate reader (BMG, Cary, NC). The protein level was calculated using the standard curve of standard samples.

### Integrative analysis with published data sets

Full list of published data sets used for comparative analyses is available in **Extended Data Table 12**. For microarray data sets (GSE93649, GSE78829 and GSE12538), CEL files were retrieved from Gene Expression Omnibus (GEO) database. Gene expression signals across samples were normalized with RMA algorithm, and differential expression analysis was performed with R package limma^126^. For RNA-seq data sets (GSE120705), sequencing reads were retrieved from Sequence Read Archive (SRA) database with SRA Toolkit 2.10.7. After the removal of adapter contamination and low-quality reads with Trim Galore v0.6.5 (https://github.com/FelixKrueger/TrimGalore), gene expression quantification was performed with Salmon 1.2.1^137^. Differential expression analysis was performed with tximeta 1.6.2 and DESeq2 1.28.0^138, 139^. For single cell RNA-seq data sets (GSE70657 and GSE59114), matrix of gene counts was obtained from GEO database. Normalization and differential expression were then performed with DESeq2 1.28.0^139^.

For chromatin immunoprecipitation sequencing (ChIP-seq) data sets (GSE49847, GSE69671, GSE34483, GSE39237, GSE16256, GSE43103, GSE59636, GSE47082, GSE43007, GSE47765, GSE76055, and GSE22075), binding peaks and gene targets were retrieved from Cistrome Data Browser. Only genes with regulatory potential score > 0.5 were used as targets for further analysis. For data sets not available in Cistrome Data Browser (GSE120705), sequencing reads were retrieved from SRA database with SRA Toolkit. Following the analytical pipeline used in Cistrome, ChiLin workflow was used for peaking calling and gene target prediction^140^. Briefly, after the removal of adapter contamination and low-quality reads with Trim Galore, genome alignment (mm10) was performed with BWA 0.7.17^141^. Unambiguously mapped sequencing reads were subjected to peak calling with MACS2 2.1.1^142^. Thereafter, target gene analysis was performed with BETA 1.0.7 to examine the regulatory potential for each RefSeq genes^143^. Binding targets with regulatory potential score > 0.5 were used as targets for further analysis. For comparative analyses performed with the data sets, the significance *p*-values for the overlapping genes were calculated by hypergeometric test with built-in phyper function of R.

### Statistical analysis

All results were expressed as the mean values ± s.e.m. unless otherwise noted. Prism8 (www.graphpad.com) was used for statistical analyses with Student’s *t*-tests. Unless otherwise specified, all statistical tests were two-sided, and analyses for significant differences between two groups of paired and unpaired samples were conducted using paired and unpaired Student’s *t*-test, respectively. Two-sample Kolmogorov-Smirnov tests were performed with built-in *ks.test* function of R, to compare the difference of frequency distribution of *Tfrc* transcript per cell under different conditions. Hypergeometric tests were performed with built-in *phyper* function of R, to assess the enrichment of IRE-containing genes and the significance of gene set overlapping. Differential expression analysis with microarray data was performed with Limma Bioconductor package^144^. Differential expression analysis with RNA-seq data was performed with DESeq2^139^. Enrichment analyses of pathways, and upstream regulators were conducted using NetworkAnalyst^145^, and Ingenuity Pathway Analysis (IPA). Enrichment analyses of gene sets were performed with GSEA for scRNA-seq data of iron chelator-treated HSC (*GSEAPreranked* analysis) and microarray data of *Fbxl5*-knockout HSC. Pre-ranked gene list for iron chelator-treated HSC were genes expressed in >10% of all the single cells in the scRNA-seq data, and ranked by the fold changes in iron chelator treatment compared to control. Signature score in single HSC was calculated with *AddModuleScore* module of *Seurat* package using the expression matrix of scRNA-seq. Estimation of stem cell frequency and significant differences in stem cell frequency between different groups in LTC-IC assays were estimated with ELDA method in the R package statmod^146^.

### Data availability

Metabolomic data collected have been included in **Extended Data Tables 6-9** in the Supplementary Information File. Transcriptomic data was deposited in GEO with accession numbers GSE157817 (Microarray gene expression profiling of mouse HSC treated with iron chelator deferoxamine (DFO)) and GSE157821 (Single cell RNA-seq of mouse HSC treated with iron chelator deferoxamine (DFO)).

## Acknowledgements

We thank the members of the Will lab for feedback on and discussions of the study, and Amit Verma for feedback on the manuscript; Sofiya Milman, Derek Huffman and Nir Barzilai for very helpful discussions, access to aged mice and *Longenity* cohort data. We further thank Daqian Sun and Swathi-Rao Narayanagari from the Stem Cell Isolation and Xenotransplantation Core and Jinghang Zhang at the Einstein Cell Sorting Core facilities for technical assistance with FACS analysis and cell sorting; David Reynolds from the Einstein Genomics Core facility for single cell capture for RNA-seq, Yunping Qiu and Xueliang Du at the Einstein-Mt Sinai Diabetes Center Stable Isotope & Metabolomics Core, the Einstein Analytical Imaging Facility, the Einstein Institute for Animal Studies, and Research Facilities Coordinator Peter Schultes for expert support and technical assistance. We also thank Justin Wheat and Goichi Tatsumi for intellectual and technical support for the smRNA FISH experiments. This study was supported by grants from the National Institutes of Health P30CA013330 (core support grant), DK105134 and CA230756 (to B.W.), as well as investigator-initiated research projects sponsored by GlaxoSmithKline and Novartis Pharmaceuticals (to B.W.), and a Pershing Square Sohn Prize for Young Investigators in Cancer Research (to B.W.). Y.R.K. is supported by a T32 Training grant in Aging Research (AG023475; PI: Barzilai) and M.M.A. is supported by the Einstein Training Program in Stem Cell Research (NYSTEM C30292GG; PI: Frenette). R.K. is the recipient of a Career Development Program Postdoctoral Fellowship from the Leukemia & Lymphoma Society (LLS).

## Author contributions

Y.R.K. and J.C. designed, performed, analysed and interpreted the cell functional, molecular, imaging and FACS-based assays, designed and conducted *in silico* data mining, and wrote the manuscript; R.K. established and performed all smRNA FISH analyses; R.K. analysed, and R.K. and U.S. interpreted the smRNA FISH data.; M.T. assisted with primary cell culture model set-up and data collection; Y.M. performed Western blot analyses; M.M.A. performed the *Ascl4* KD model set up and validation, and *Ascl4* KD LTC-IC data collection; A.Z., V.T. and M.T. performed mouse colony management, dissection and harvest of hematopoietic cells from the bone marrow of animals; S.S.L. and J.H. performed and collected data for mass spectrometry-based proteomic and metabolomic analyses; S.S. and A.D’A. coordinated and interpreted mass spectrometry-based proteomic and metabolomic analyses; B.W. designed and coordinated the study, contributed to designing the experiments and interpreting the data, and wrote the manuscript.

## Additional information

**Supplementary Information** is available for this paper.

**Correspondence and requests for materials** should be addressed to Yun-Ruei Kao, Ph.D. or Britta Will, Ph.D.

## Extended data figures

**Extended Data Figure 1.**
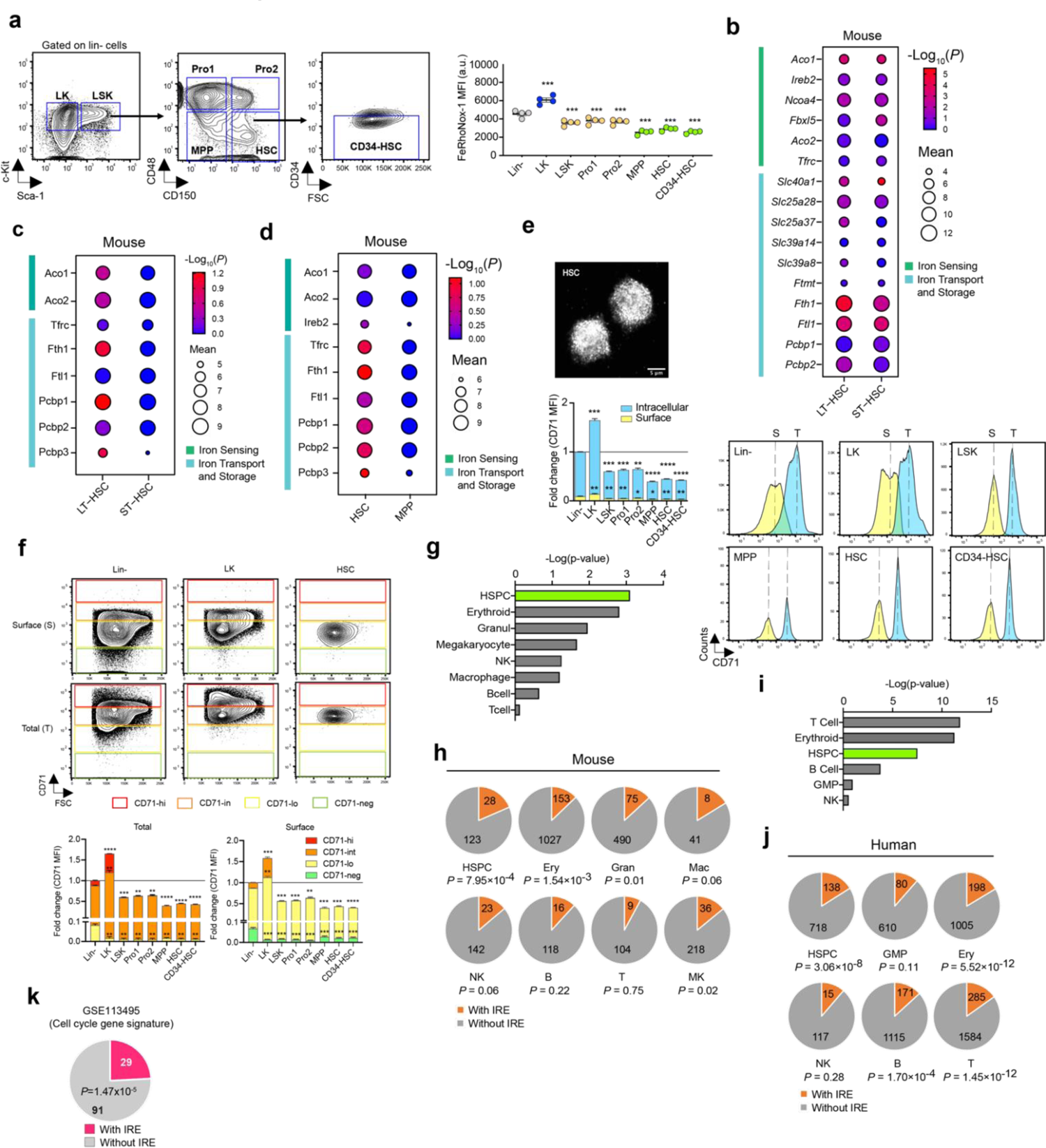
Iron-dependent regulation of hematopoietic stem cell defining gene expression. **a,** Quantification of the intracellular labile iron pool (LIP) using FeRhoNox-1 and FACS analysis. Upper panel: Gating strategy for hematopoietic stem and progenitor cell (HPSC) populations; lower graph: Quantification of FeRhoNox-1 mean fluorescence intensity (MFI in arbitrary units (a.u.)). n = 4. **b,** mRNA expression of iron homeostasis-regulatory genes in HSC compared to progenitor cells (PMID: 18371395). Expression of genes with regulatory functions in iron sensing or transport and storage were examined. The normalized log_2_ expression values of relevant genes in LT-HSC or ST-HSC were compared to megakaryocyte-erythroid progenitor (MEP), n = 2-5. **c,d,** Protein expression of iron homeostasis-regulatory genes in murine HSPC (**c**: PMID25158935; **d**: PMID26299573). The levels of proteins with regulatory functions in iron sensing or transport and storage were examined. The log_10_ peptide intensities across HSC and progenitor cells were compared. n = 3. Significance p-value were calculated using unpaired Student’s *t*-test by comparing LT-HSC versus ST-HSC (**c**), or HSC versus MPP (**d**). **e,** CD71 detection in purified HSC by immunofluorescence and confocal microscopy and quantification of cell surface and total CD71 content in HSPC populations by FACS analysis. The fold changes of CD71 MFI in different hematopoietic stem and progenitor cells compared to Lin-population are shown, n = 4. **f,** CD71 cell surface (S) and total (T) protein abundance in phenotypically defined hematopoietic stem and progenitor cell populations. Representative FACS plots for Lineage-negative (Lin^-^), Lin^-^ cKit^+^Sca-1^-^ (LK) and Lin^-^ckit-Sca-1^+^CD48^-^CD150^+^ (HSC) cells (upper) and quantification (bottom). CD71-negative (CD71-neg), CD71-low (CD71-lo), CD71-intermediate (CD71-int), and CD71-high (CD71-hi) fractions were shown. Quantification presented as fold change of CD71 mean fluorescence intensity (MFI) in each cell populations compared to Lin^-^ cells for total CD71 protein levels (left graph) and cell surface presentation (right graph), n = 4. **g,h,** IRE enrichment analysis in murine hematopoietic cell type-specific gene expression signatures previously defined (**g**, GSE77098). **h,** Pie charts show the number of analyzed transcripts containing at least one IRE motif (*With IRE*, orange) or lacking it (*Without IRE*, gray). Hypergeometric test was used to calculate the significance *p*-value of IRE enrichment. **i,j,** IRE motif enrichment analysis in human hematopoietic cell type-specific gene expression signatures previously defined (GSE24759). **k** IRE enrichment analysis of genes associated with HSC cycling as previously defined (**k,** PMID29915358/GSE113495). If not specified differently, data are mean ± SEM. (**a, e, f**). Significance *p*-values, indicated as \**p* < 0.05, \*\**p* < 0.01, \*\*\**p* < 0.001, \*\*\*\**p* < 0.0001, were calculated by Student’s *t*-test (paired: **a, e, f;** unpaired: **b, c, d**), or Hypergeometric test (**g-j**).

**Extended Data Figure 2.**
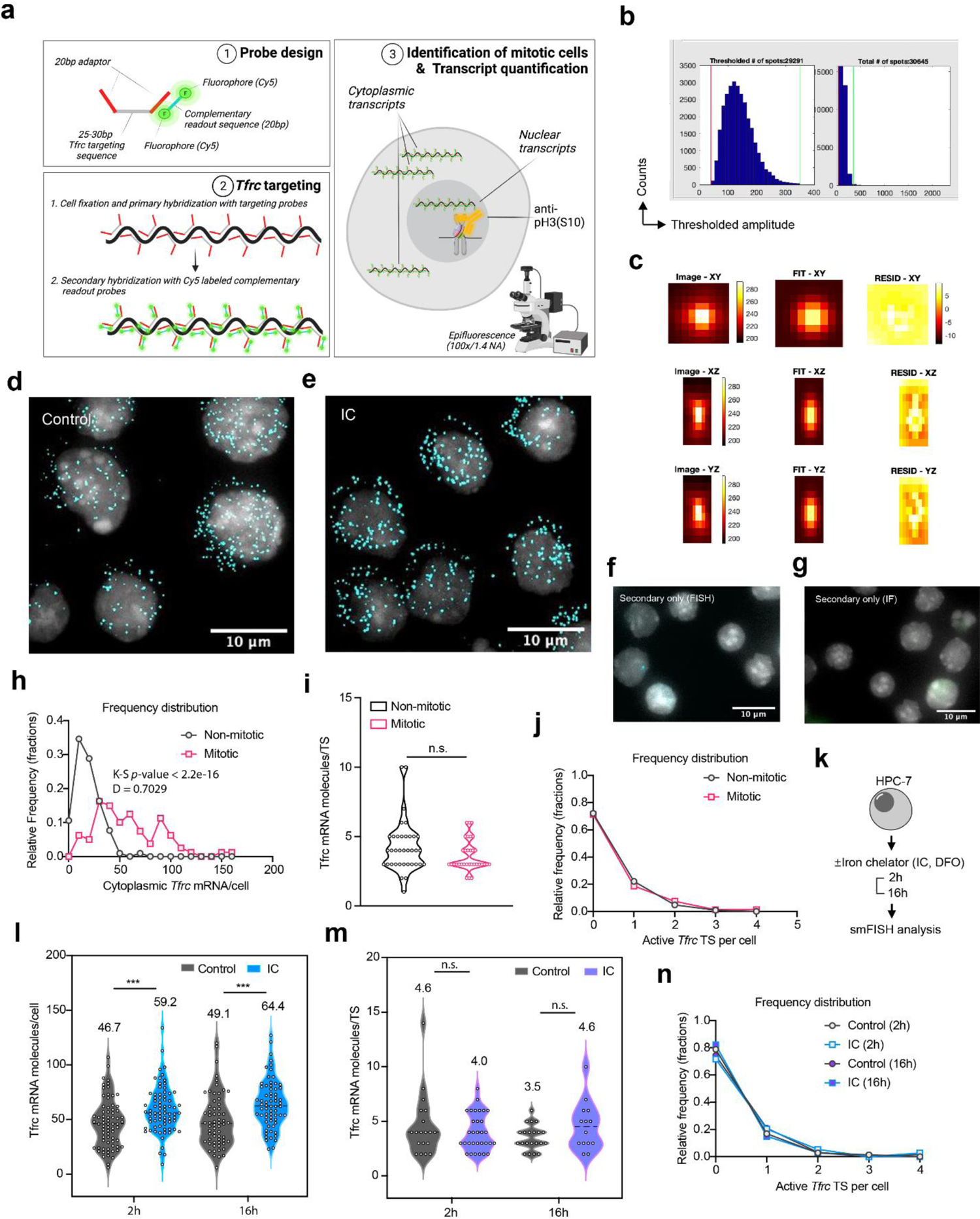
*Tfrc* single mRNA molecule analysis in HSPC. **a**, Scheme showing sequential smRNA FISH procedure for *Tfrc*. **b,** Computational thresholding of amplitude of *Tfrc* mRNA. **c**, Fit of averaged image of *Tfrc* mRNA with 3D Gaussian distribution. First row: maximum intensity projections in XY; second row: maximum intensity projections in XZ. The first column shows the image, the second column the best fit, and the third column the absolute residuals. **d-g**, Representative filtered and overlaid images of cells stained by smRNA FISH for *Tfrc* (Cy5, cyan pseudocolor), Control (**d**, HPC-7) vs. IC-treated (**e**, HPC-7), as well as secondary probe only controls for smRNA FISH (**f,** RO2-Cy5, Lin-cKit+ (LK) cells) and immunofluorescence (**g**, anti-rabbit AlexaFluor 488, LK cells) are shown. Nucleus is stained with DAPI (gray pseudocolor). Scale bars, 10 μm. **h,** Histogram showing frequency distribution of cytoplasmic *Tfrc* mRNA per cell in non-mitotic and mitotic HSC (HPC-7). **i,** Violin plot showing *Tfrc* nascent mRNA molecules per transcription site (TS) in non-mitotic (n=36) and mitotic HSC (n=34). **j,** Histogram showing frequency distribution of active *Tfrc* transcription sites (TS) per cell in non-mitotic and mitotic HSC. **k**, Schematic showing experimental flow for HPC-7 cells exposed to either control (H_2_O) or IC (DFO) for 2 or 16hrs followed by *Tfrc* smRNA FISH analysis. **l**, Violin plot showing total *Tfrc* mRNA molecules per cell. For 2h, n=71 (Control) and n=74 (IC); for 16h, n=62 (control) and n=66 (IC). **m**, Violin plot showing *Tfrc* nascent mRNA molecules per transcription site (TS). For 2h, n=19 (Control) and n=31 (IC); for 16h, n=21 (control) and n=14 (IC). **n**, Histogram showing frequency distribution of active *Tfrc* transcription sites (TS) per cell in non-mitotic and mitotic cells. Significance p-values, indicated as *p < 0.05, **p < 0.01, ***p < 0.001, ****p < 0.0001, were calculated using Student’s t-test (unpaired: **i, l and m**) and two-sample Kolmogorov-Smirnov tests (**h**).

**Extended Data Figure 3.**
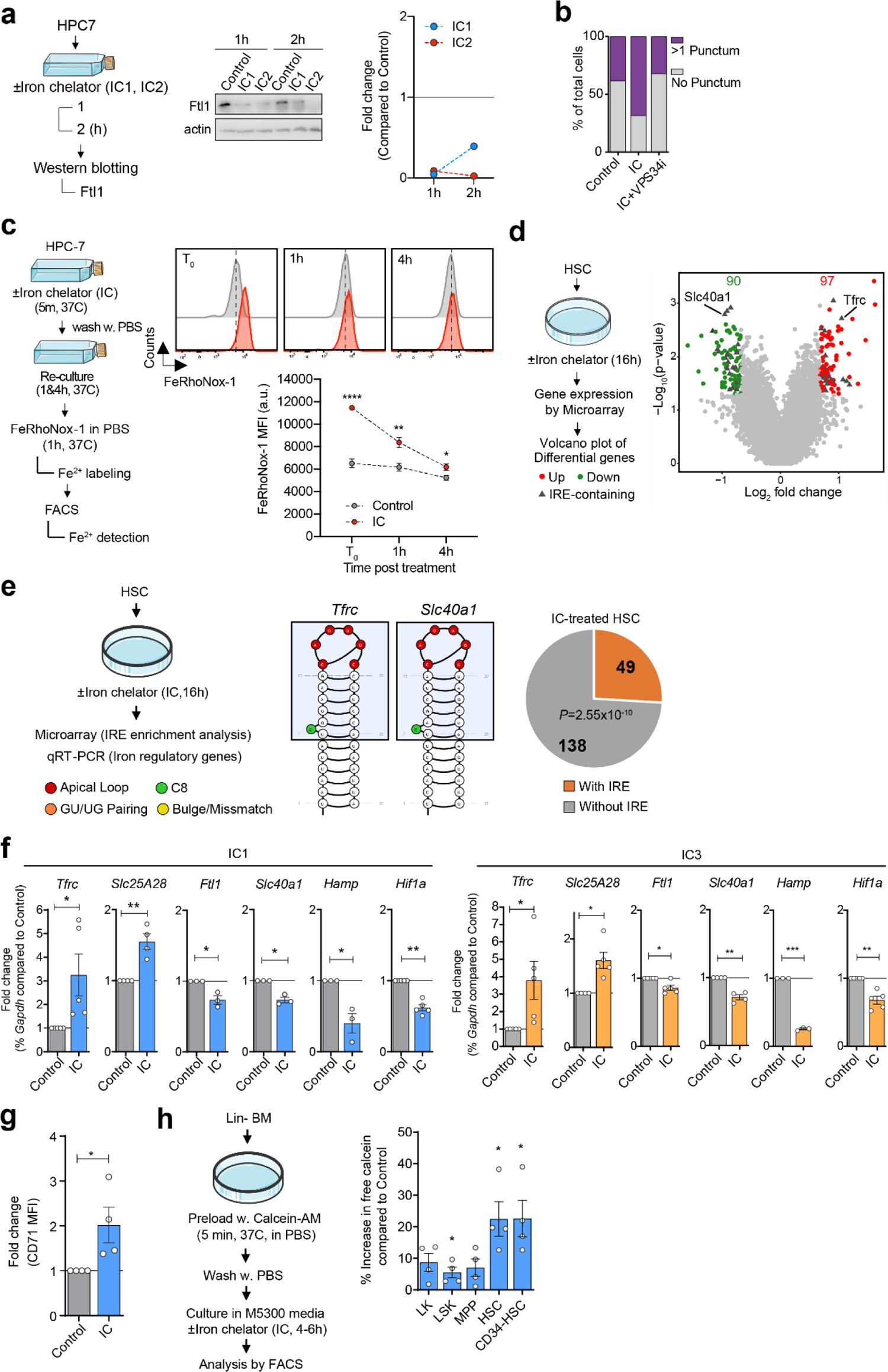
Iron chelator-mediated induction of the limited-iron response in HSPC. **a,** Ftl1 protein quantification in HPC-7 cells at 1 and 2hrs after IC (IC1: DFO, IC2: EP) exposure. Western blot images (middle) and quantification (right). **b,** Immunofluorescence and confocal microscopy analysis of Ncoa4-dependent ferritinophagy in HSC after IC treatment for 48hrs. The number of Noca4 puncta in each single HSC was enumerated, and the frequency of cells with at least one Ncoa4 punctum is shown. **c,** Quantification of alteration in intracellular Fe^2+^ content after IC (EP) treatment of HPC-7 cells using FeRhoNox-1 and FACS analysis. Scheme (left) and representative FACS plots (right) at different time points. n = 6; 2 independent experiments. **d,** Left: Scheme of microarray analysis to identify expression changes in HSC after 16hrs IC (DFO) exposure compared to mock treatment controls. Right: Volcano plot showing the distribution of expression changes. Significantly down- or up-regulated genes are highlighted, and IRE-containing genes are denoted with triangles, includes *Tfrc* and *Slc40a1*. **e,** Gene expression changes in purified HSC 16hrs after IC exposure (IC1: DFO) compared to mock treatment controls by microarray identifies differential levels of IRE-containing transcripts (pie chart) and includes *Tfrc* and *Slc40a1* (IRE structures shown as example). **f,** Alterations in the mRNA expression of the iron regulatory pathway assessed by qRT-PCR (upper IC1: DFO; lower IC3: DFX), n = 3-5. **g,** Cell surface CD71 presentation on HSC 4-6hrs upon iron chelator (DFO) exposure by FACS analysis, n = 4. **h,** Quantification of the labile iron pool in HSPC populations. Scheme shows cells preloaded with Calcein-AM exposed to IC or vehicle treatment in growth medium. Data represents the increase in Calcein-AM mean fluorescence intensity (MFI) upon IC treatment (% vehicle control). n = 4. If not specified differently, data are mean ± SEM. (**c, f, g, h**). Significance *p*-values, indicated as \**p* < 0.05, \*\**p* < 0.01, \*\*\**p* < 0.001, calculated by Student’s *t*-test (paired: **f, g, h;** unpaired: **c**), or Hypergeometric test (**e**).

**Extended Data Figure 4.**
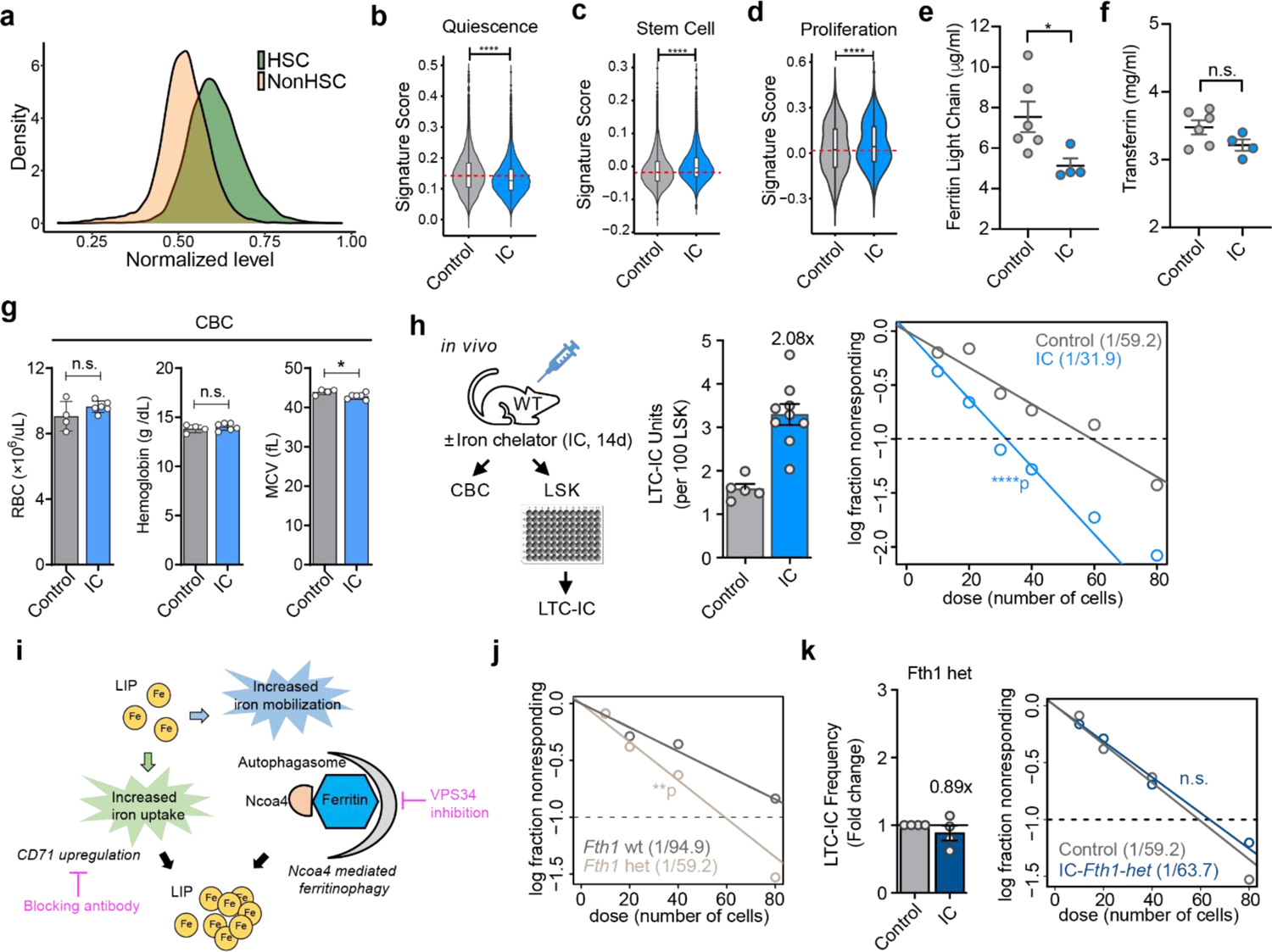
Activation of the limited-iron response increases HSPC. **a,** Distribution of union HSC signature expression in phenotypic HSC (green) and non-HSC (orange). Average expression of the signatures in each of the single cells were calculated. Density plot shows the Kernel density estimate of the distribution of HSC signature expression in phenotypic HSC versus non-HSC. **b,c,** Comparison of the expression of IC-treated HSC versus control HSC in scRNA sequencing, with gene signatures associated with HSC quiescence (**b,** GSE108155) and stemness (**c**). Lists of the gene signatures are available in **Extended Data Table 1**. Signature scores were calculated with *Seurat* package. **d,** Comparison of the expression of IC-treated HSC versus control HSC in scRNA sequencing with the cycling signatures (GSE113495). **e,f,** Quantification of protein levels of Ferritin light chain (**e**) and transferrin (**f**) in the serum of mice exposed to *in vivo* treatment of IC (DFO) or vehicle control for 2 weeks. **g,** Complete blood count (CBC) analysis on peripheral blood of mice after *in vivo* iron chelator (*IC*) or mock control (*control*) treatment. Quantification of red blood cells (RBC), hemoglobin, and mean corpuscular volume (MCV). n = 4-6. **h,** Quantification of functional HSC after *in vivo* iron chelator (IC: DFO) treatment using LTC-IC assay. Left: bar plot shows the LTC-IC frequencies across individual mice in the group of IC or control. n = 5 (control) and n = 9 (IC). Estimated group LTC-IC frequencies by ELDA (right); ELDA-calculated LTC-IC frequencies for both groups in parentheses in ELDA plot (right). **i,** Experimental strategy for pharmacological inhibition of iron import and intracellular mobilization. **j, k,** LTC-IC assay to quantify HSC frequency in LSK cells without and with heterozygous deletion of *Fth1* (**i**), and HSC frequency in *Fth1*-*het* LSK following mock or IC (DFO) treatment (**j**), n = 4. If not specified differently, data are mean ± SEM. (**e, f, g; h, k** (bar graph)). Significance *p*-values, indicated as \**p* < 0.05, \*\**p* < 0.01, \*\*\**p* < 0.001, \*\*\*\**p* < 0.0001, calculated by Student’s *t*-test (paired: **g;** unpaired: **b-f**), or Poisson statistics: **h, j, k** (ELDA analysis).

**Extended Data Figure 5.**
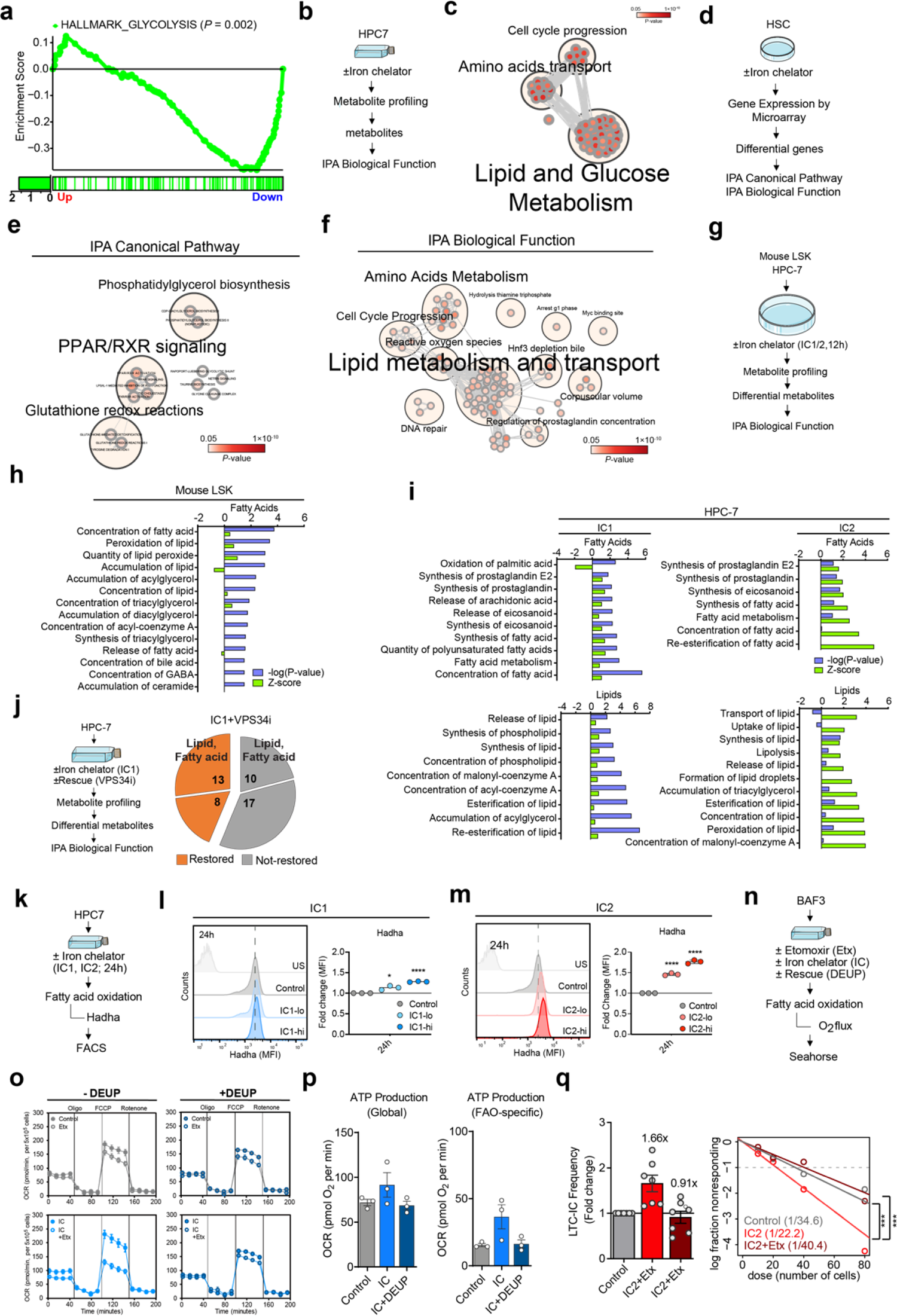
Activation of the limited iron response stimulates fatty acid oxidation in HSPC. **a.** *GSEAPreranked* analyses of scRNA-seq data using MSigDB gene set HALLMARK_GLYCOLYSIS. **b,c,** Following IC (DFO) treatment, HPC-7 cells were subjected to metabolite profiling (**b**). Differential metabolites were subjected to pathway analysis with IPA. **c,** Pathway network analysis of differential metabolites in HPC-7 cells after 12-hours IC exposure, n = 4. **d-f, d,** Scheme for assessing the transcriptional alterations triggered by iron chelation. Sorted HSC were subjected to *ex vivo* treatment with DFO for 16 hours, followed by microarray analysis. IPA pathway analysis was performed with differentially expressed genes (DEG; average fold change > 1.6 and p < 0.05; n=4). **e,f,** Pathway network analysis of DEG by IPA canonical pathway analysis (**f**), and biological functions (**f**), and significantly enriched terms (*p* < 0.05) were subjected to *EnrichmentMap* for network visualization, and clustered with *AutoAnnotate*. **g-i,** Metabolite profiling of mouse Lin^-^Sca-1^+^cKit^+^ (LSK) (**h**) and HPC-7 cells (**i**) following the *ex vivo* treatment with iron chelators for 12hrs (**g**). **h,i**, IPA biological function analysis performed with metabolites showing significant changes upon IC (DFO) treatment (with an average fold change > 1.3) compared to mock treatment controls. Pathways related to fatty acid and lipid metabolism are shown for LSK (**h**, n = 2), HPC-7 cells (**i**, n = 4). **j,** VPS34i reverses differential abundance upon IC (DFO) exposure of almost half of the metabolites, including a significant number of metabolites belonging to fatty acid and lipid metabolic pathways as indicated. Restored metabolites were defined as those with average fold change > 1.25 (IC1 versus IC1+VPS34i; n = 3). **k-m, k,** Hadha staining in HPC-7 cells exposed to iron chelators (IC1-lo: DFO 5μM, IC1-hi: DFO 10μM, or IC2-lo: EP 3µg/ml, IC2-hi: EP 10µg/ml) or respective mock treatment controls for 24hrs. **l,m,** Quantification of Hadha MFI upon the treatment of IC1 (**l**) or IC2 (**m**), representative histograms and fold changes of Hadha mean fluorescence intensity (MFI) values compared to mock controls are shown. n = 3. **n,o, n**, Assessment of fatty acid metabolism in hematopoietic progenitor cell line, Ba/F3, by Seahorse analysis detecting total OXPHOS (control) and β-oxidation (after CPT-1 inhibition with etomoxir (Etx)) specific oxygen consumption rates (OCR) in the presence or absence of IC (DFO), without (left) or with (right) inhibition on lipolysis using diethylumbelliferyl phosphate (DEUP). **o,** representative analysis plots. **p,** Total (left) and FAO-specific (right) mitochondrial ATP production measured by Seahorse analysis. n = 3. **q,** LTC-IC assay to quantify functional HSC upon or IC2 (EP) treatment alone or in combination with pharmacological inhibition of fatty acid import to mitochondria using etomoxir (Etx). Fold changes of long-term culture-initiating cell (LTC-IC) frequencies compared to controls across individual mice (bar graph). In the ELDA plot, average stem cell frequency is shown for each treatment group. n = 7. If not specified differently, data are mean ± SEM. (**p, q**). Significance *p*-values, indicated as \**p* < 0.05, \*\**p* < 0.01, \*\*\**p* < 0.001, \*\*\*\**p* < 0.0001, were calculated by Student’s *t*-test (paired: **l, m**).

**Extended Data Figure 6.**
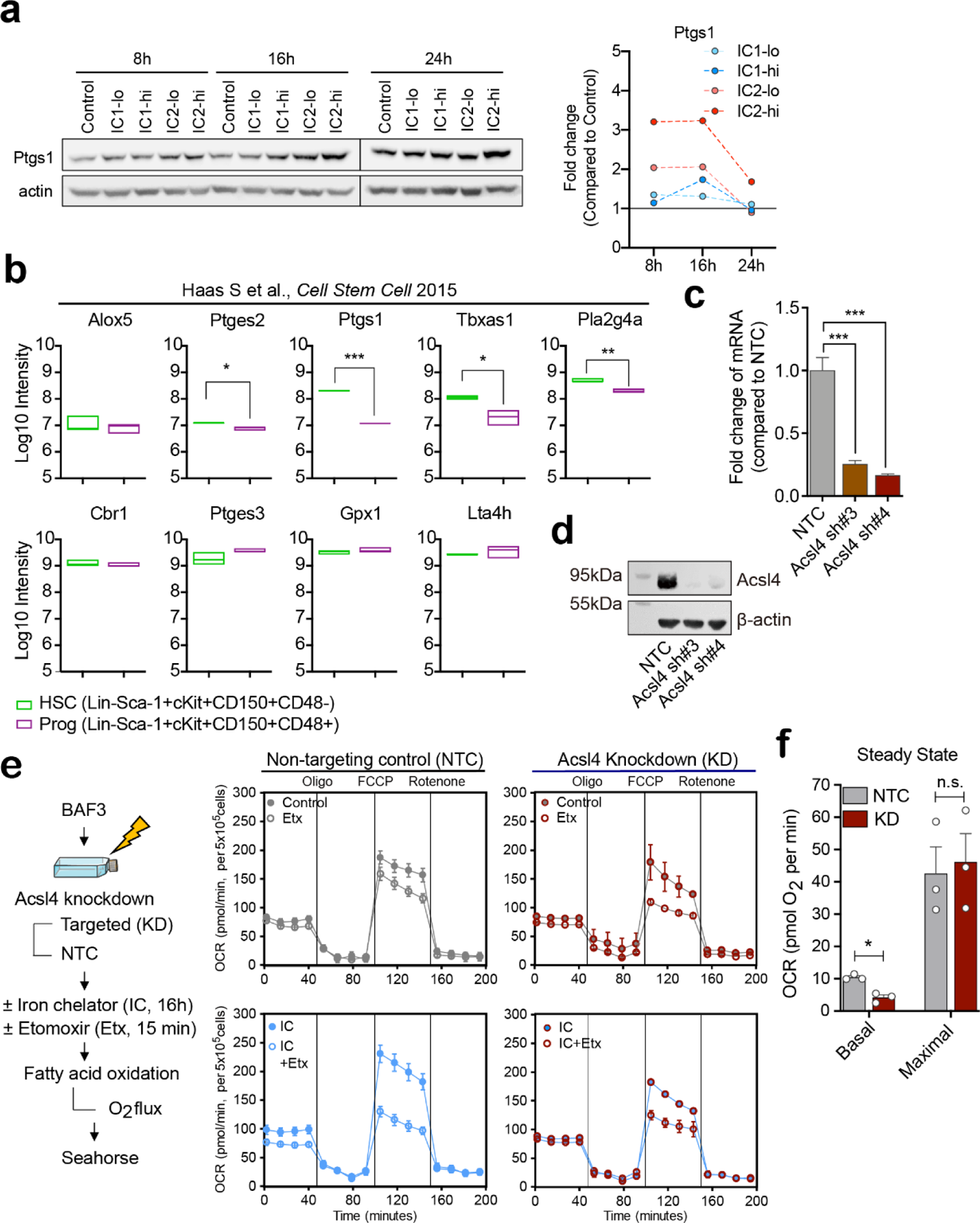
β-oxidation of arachidonic acid increases HSPC pool upon limited iron response activation. **a,** Protein expression of Ptgs1 in HPC-7 cells upon treatment with IC1 (DFO), IC2 (EP) or in mock treatment controls at 8, 16 and 24hrs after IC exposure. Representative images (left panels) and fold changes of actin-normalized protein levels by image analysis compared to vehicle control are shown (right graphs). **b,** Protein expression of genes involved in arachidonic acid metabolism in HSC versus progenitors (*Prog*). Peptide intensities across HSC and progenitor cells^62^ were compared (PMID: 26299573), n = 3. **c,d,** Genetic inhibition of *Acyl-CoA Synthetase Long Chain Family Member 4* (*Acsl4*) by RNAi in primary cKit+ bone marrow mononuclear cells using two independent shRNA targeting *Acsl4* (*Acsl4 sh#3*, *Acsl4 sh#4*) or a non-targeting control (*NTC*). Quantification of *Acsl4* mRNA expression (**c**) and protein levels (**d**) in successfully transduced (GFP^+^) cells 48hrs after lentiviral transduction, n = 3. **e,** Assessment of fatty acid oxidation rates in BA/F3 cells transduced with non-targeting control (NTC) or shRNA targeting *Acsl4* (KD) after IC (DFO) treatment. Experimental scheme and representative OCR plots for NTC (left plots)) and *Acsl4* knockdown (right plots) cells. **f,** Assessment of fatty acid oxidation rates in BA/F3 cells transduced with non-targeting control (NTC) or shRNA targeting *Acsl4* (KD) under steady state. Basal and maximal fatty acid-specific OCRs are shown, n = 3. If not specified differently, data are mean ± SEM (**b, c, f**). *p < 0.05, **p < 0.01, ***p < 0.001, *n.s*. not significant, calculated by Student’s *t*-test (unpaired: **b, c**; paired: **f**)

**Extended Data Figure 7.**
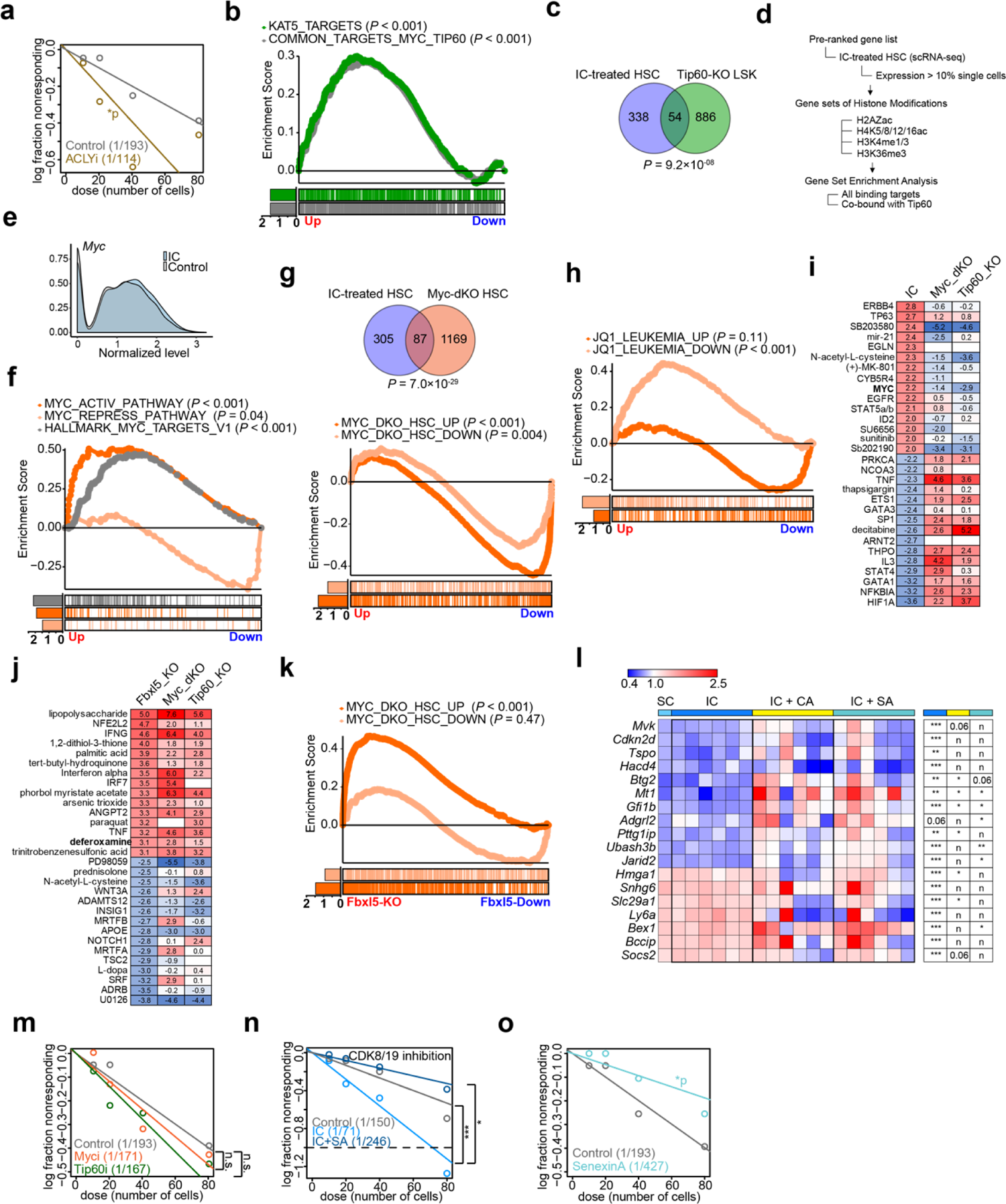
Increased HSC self-renewal associated gene expression upon iron limitation response activation. **a,** Quantification of HSC by LTC-IC assay after the single treatment with inhibitor of ATP citrate lyase (ACLY). **b,** *GSEAPreranked* analyses of scRNA-seq data using MSigDB gene set KAT5_TARGET_GENES (human cells), and common binding targets of Tip60 and Myc. Common binding targets of Tip60 and Myc were obtained by comparing the ChIP-seq of Tip60 in ESC and Myc in CH12 and MEL cells published previously (GSE49847). Only genes bound by Myc in both CH12 and MEL cells were considered as Myc target in this analysis. **c,** Overlap of DEG in IC-treated HSC by scRNA sequencing with those in Tip60-KO LSK cells. Number of unique and shared DEG are shown. Significance of overlap was calculated using Hypergeometric tests. **d,** Analysis strategy for **Figure 3c**. **e-h,** Transcriptomic analysis of purified HSC 48hrs after IC (DFO) exposure or mock treatment using singe cell RNA-seq (scRNA-seq) analysis. **e,** Myc expression in IC treated HSC (blue) and vehicle controls (grey). **f,** *GSEAPreranked* analysis using MSigDB collection gene sets of validated targets upon c-Myc transcriptional activation (PID_MYC_ACTIV_PATHWAY) and repression (PID_MYC_REPRESS_PATHWAY), as well as HALLMARK_MYC_TARGETS_V1. **g,** Upper panel: overlap of differentially expressed genes (DEG) in IC-treated HSC (vs. vehicle control) and HSC lacking *c-Myc* and *n-Myc* (Myc-dKO)^147^. Number of unique and shared DEG are shown. Significance of overlap was calculated using Hypergeometric tests. Lower panel: *GSEAPreranked* analyses using DEG in Myc-dKO versus wildtype. **h,** *GSEAPreranked* analyses with gene set of DEG upon JQ1-treatemt in K562 and MV4-11 cells^148^. **i,** IPA upstream regulator analysis comparing DEG in IC-treated HSC, Myc-dKO HSC^147^, and Tip60-KO LSK^72^. Z-scores of top 15 activated and inhibited (ranked by Z-score) upstream regulators and corresponding Z-scores in Myc-dKO HSC and Tip60-KO LSK are shown. **j,** IPA upstream regulator analyses of DEG (absolute fold change > 1.5, *p*-value < 0.05) in *Fbxl5*-deficient HSC (Fbxl5-KO)^43^ were compared to gene expression alterations seen in *Myc*-dKO HSC^147^ and *Tip60*-KO LSK (vs. wildtype controls). Z-scores of top 15 activated (ranked by Z-score) and inhibited upstream regulators in *Fbxl5*-KO HSC and corresponding Z-scores in *Myc*-dKO HSC and *Tip60*-KO LSK are shown. **k,** GSEA analysis of expression profile of *Fbxl5*-KO HSC was performed with DEG in *Myc*-dKO HSC versus wildtype controls; significantly up-regulated DEG enrichment analysis (orange) and down-regulated genes (light orange). **l,** Expression changes of target genes in HSC after IC treatment alone or in combination with CDK8/CDK19 inhibitors (CA: Cortistatin A; SA: Senexin A) by Fluidigm analysis. Fold changes of genes across treatment groups from Fluidigm and scRNA-seq (SC) are shown. Significance of differential expression shown (right panel) were estimated by paired Student’s *t*-test comparing the Δ^Ct^ in IC versus control, co-treatment versus IC. *p < 0.05, **p < 0.01, ***p < 0.001, *n*, not significant. **m-o,** Quantification of HSC by LTC-IC assay after the single treatment with inhibitors of Tip60 or Myc (**m**), CDK8/CDK19 (SA: Senexin A, **o**) or in combination with IC (**n**). Average LTC-IC frequencies are shown in parentheses. Significance tested using ELDA. Significance *p*-values, indicated as \**p* < 0.05, *n.s.* not significant, calculated by Poisson statistics.

**Extended Data Figure 8.**
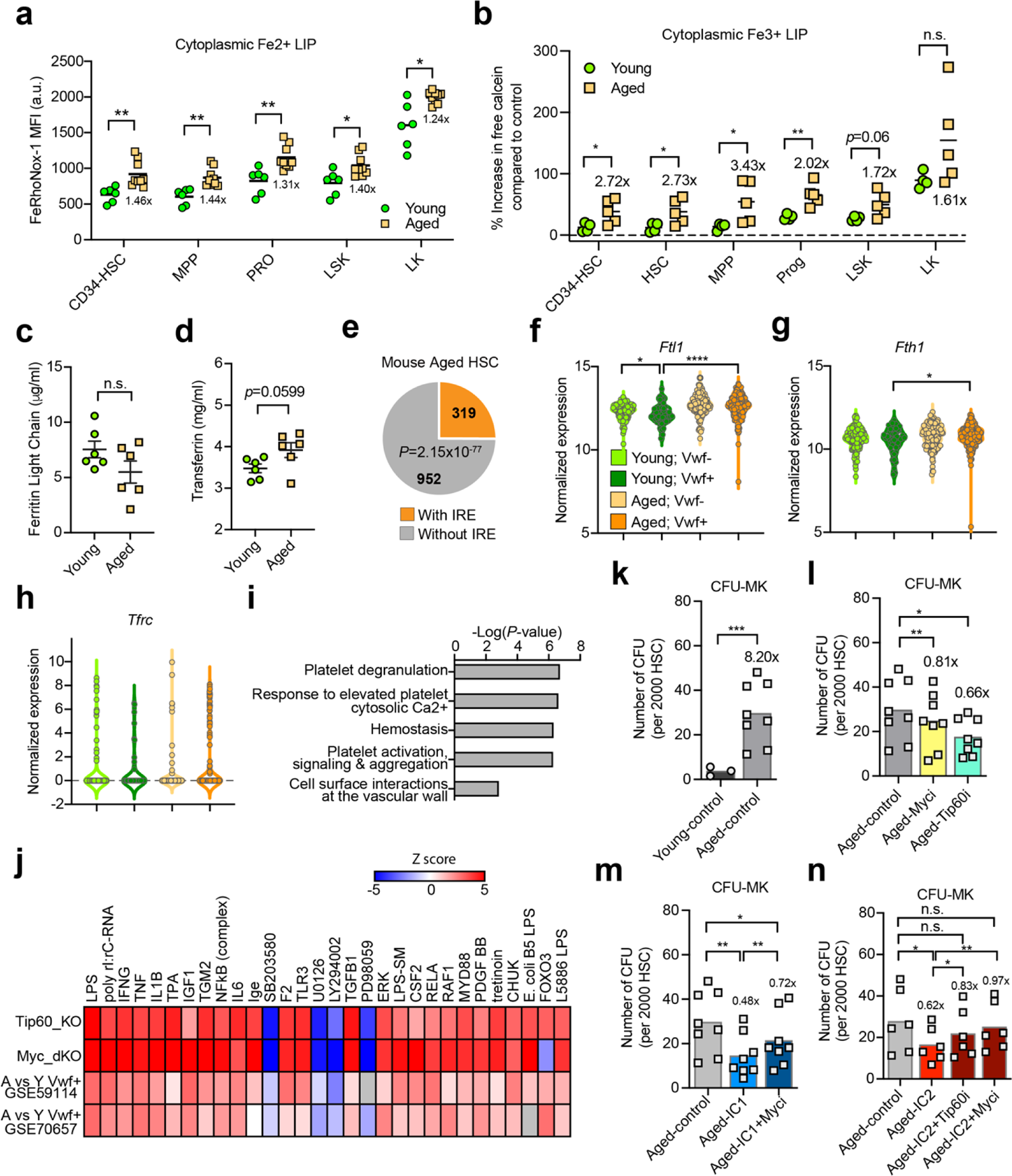
Cytoplasmic iron loading compromises function of aged HPSC. **a,b,** Increased labile iron pool (LIP) in aged hematopoietic stem and progenitors compared to young mice. Cytoplasmic Fe^2+^ (**a**) or Fe^3+^ (**b**) LIP were measured using FeRhoNox-1 and Calcein-AM staining, respectively. Quantification of FeRhoNox-1 and Calcein-AM mean fluorescence intensity (MFI in arbitrary units (a.u.)) by FACS analysis are shown. n = 4-9. **c,d,** Quantification of protein levels of Ferritin light chain (**c**) and transferrin (**d**) in the serum of aged versus young mice. **e,** IRE enrichment analysis with DEG of murine aged HSC compared to young HSC in GSE47817. Significance of enrichment was calculated by hypergeometric test. **f-h,** Comparison of the expression of *Ftl1* (**f**), *Fth1* (**g**), and *Tfrc* (**h**) in HSC isolated from young and aged mice. Single cell RNA sequencing data of aged and young HSC was obtained from previous study (GSE59114); HSC with Von Willebrand Factor (*Vwf*) expression >3 were defined as Vwf+, The expressions of *Ftl1*, *Fth1*, and *Tfrc* were compared across different groups. **i**, Pathway analysis of genes in the leading edge of GSEA shown in **Fig. 4g**. **j,** IPA upstream regulator analysis comparing DEG in Myc-dKO HSC^147^, Tip60-KO LSK^72^, as well as DEG in aged versus young Vwf+ HSC from previous studies (GSE59114 and GSE70657). Z-scores of top 15 activated and inhibited (ranked by Z-score) upstream regulators and their corresponding Z-scores are shown. **k-n,** Quantification of megakaryocyte colony-forming units in aged HSC compared to young HSC at baseline level (**k**). CFU-Mk derived from aged HSC in response to Tip60 or Myc inhibitors alone (**l**), or in combination with iron chelators (**m, n**). Quantification of CFU-Mk colonies across different conditions was normalized to 2000 HSC plated. n = 3-8. If not specified differently, significance p-values, indicated as *p < 0.05, **p < 0.01, ***p < 0.001, ****p < 0.0001, were calculated using Student’s t-test (unpaired: **a**-**d, f-h, j**, and paired: **k-m**).

**Extended Data Figure 9.**
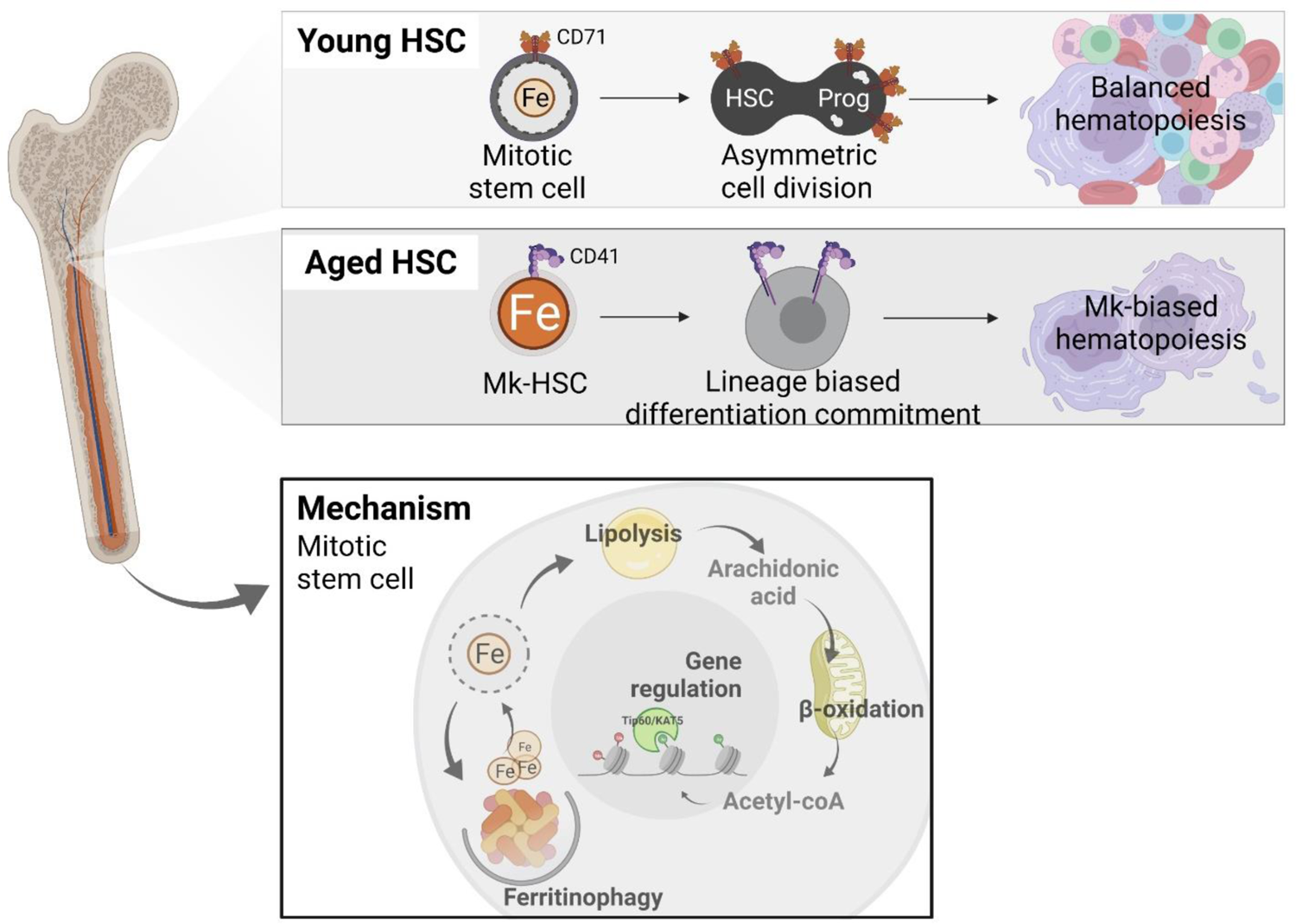
Proposed cell functional and molecular model.

---

**Extended Data Table 1.** List of gene signatures.

**Extended Data Table 2.** Differentially expressed genes in DFO-treated HSC by Microarray analysis.

**Extended Data Table 3.** Differentially expressed genes in DFO-treated HSC by scRNA-seq.

**Extended Data Table 4.** Pathway enrichment of differentially expressed genes in DFO-treated HSC by Microarray analysis.

**Extended Data Table 5.** Pathway enrichment of differentially expressed genes in DFO-treated HSC by scRNA-seq.

**Extended Data Table 6.** Altered metabolites in HPC7 cells with the treatment of DFO.

**Extended Data Table 7.** Altered metabolites in mouse LSK cells with the treatment of DFO.

**Extended Data Table 8.** Biological function enrichment (IPA analysis) of altered metabolites in DFO-treated HPC7 cells.

**Extended Data Table 9.** Biological function enrichment (IPA analysis) of altered metabolites in DFO-treated LSK cells.

**Extended Data Table 10.** List of antibodies used in the study.

**Extended Data Table 11.** List of oligos used in this study.

**Extended Data Table 12.** List of published data sets used for integrative analysis.

